# A Neuronally Active Bacterial Siderophore Manipulates Host Behavior and Longevity

**DOI:** 10.64898/2026.09.30.755717

**Authors:** Yifan Wu, Saul Moore, Cassandra Backes, Alan Koh, Aleksandra Zečić, Daniel W. Djamriani, Björn Heinemann, Cecile Angermann, Riju Balachandran, Tanara V. Peres, Jennifer van der Laan, Ali Ahmadi, Julian Strippel, André-Marcel Wesseling, Iliyana Kaneva, Alex Montoya, Susanne Brodesser, Helena M. Cochemé, Stephanie Kath-Schorr, Aleksandra Trifunovic, Tatjana Hildebrandt, André E.X. Brown, Filipe Cabreiro

## Abstract

Aging populations face a growing burden of neurodegenerative disease and behavioral decline. The microbiota is increasingly implicated in host physiology, highlighting an urgent need to understand how gut microbes modulate brain function and late-life health. Here, we develop a high-throughput microbe–host screening platform to systematically dissect how bacterial genes shape organismal behavior. Using the *Escherichia coli* Keio library of 3,985 distinct non-essential single-gene deletion mutants in combination with *Caenorhabditis elegans*, we quantitatively profiled 768 behavioral features per bacterial clone and uncovered a coherent module of bacterial genes involved in iron homeostasis that robustly reprogrammed host behavior. Bacterial omics analyses revealed that alterations in iron homeostasis amplify siderophore production in a TCA cycle-dependent manner, and loss of enterobactin biosynthesis abolished the bacteria-induced changes in worm movement. Transcriptional, molecular and functional studies demonstrated that bacterial enterobactin activates host neuromuscular responses to shape both lifespan and healthspan in a manner dependent on the mitochondrial unfolded protein response. Thermal proteome profiling identified mitochondrial glutaryl-CoA dehydrogenase and the electron-transfer flavoproteins A and B as enterobactin-binding targets that control mitochondrial function. Finally, genetic manipulation of neuronal mitochondrial and cytosolic reactive oxygen species via superoxide dismutase and catalase revealed that redox state modulates enterobactin-driven changes in movement and lifespan through the secretin-family GPCR neuropeptide receptor PDFR-1. Together, these findings establish microbial enterobactin as a non-canonical neurotransmitter regulating neuronal mitochondrial function, behavior and aging, defining a gut bacteria–brain axis with broad implications for maintaining late-life health.

## INTRODUCTION

Aging is accompanied by progressive declines in neuromuscular function, behavioral coordination, and resilience, reflecting intertwined deterioration of mitochondrial homeostasis and neuronal health. Increasing evidence suggests that the gut microbiota exerts a profound influence on host metabolism ^1^ and immune regulation ^2^, yet how microbial metabolites and macromolecules directly modulate nervous system function remains largely unresolved. The gut microbiota impacts behavior and influences brain development and myelination patterns through modulation of neurotransmitter production in both the gut and brain ^3^. Studies have previously reported that levels of the microbial metabolite 4-ethylphenyl sulfate (4EPS) are increased in a mouse model of atypical neurodevelopment ^4^, and that gut bacteria bioengineered to produce 4EPS alter oligodendrocyte maturation and myelination, change brain connectivity, and induce anxiety-like behavior ^5^. Further, short-chain fatty acids (SCFAs), tryptophan and indole derivatives, trimethylamine-oxide, bile acids, polyunsaturated fatty acids, and branched-chain amino acids have been linked to neuroinflammation and neurodegeneration ^6^. The complex regulation of brain function by the microbiota also involves the modulation of multiple neurotransmitter networks, including the dopaminergic, cholinergic, serotonergic, glutamatergic, and GABAergic systems ^7^.

According to the endosymbiotic theory, mitochondria originated from the engulfment of a bacterium by an ancestral host cell approximately 2 billion years ago. This shared evolutionary origin, together with the central metabolic and signaling roles of mitochondria, provides a compelling framework for understanding how the gut microbiota regulates systemic physiology ^8^. The communication between microbes and mitochondria is emerging as a key axis of host regulation, with growing evidence that bacterial metabolites fine-tune mitochondrial metabolism and dynamics, influencing host intestinal homeostasis ^8,9^, longevity ^10,11^, and responses to therapy ^12,13^. In particular, iron metabolism, which is essential for both bacterial survival and host mitochondrial activity, represents a potential axis of interspecies communication, with microbial siderophores serving as key mediators of iron acquisition. Previous reports have shown that iron chelation by enterobactin facilitates the transport of iron into the mitochondria of *C. elegans* and human cells through interaction with the alpha subunit of ATP synthase to promote mitochondrial iron homeostasis and to regulate development. in the worm host ^14,15^.

The nematode *C. elegans* provides a powerful genetic model to dissect how aging, behavior and the microbiota intersect at an organismal scale ^16^. Its short lifespan, well-annotated cell lineages and connectome allow quantitative analysis of age-associated decline in locomotion, neuromuscular function and cognitive-like behaviors under tightly controlled environmental conditions. The animal’s tractable microbiota, which is classically modelled using defined *E. coli* strains, enables precise manipulation of bacterial genes and metabolites to probe causal microbe–host relationships. Extensive toolkits for genetics, live imaging and automated behavioral tracking in 96-well plates ^17^ make it possible to link specific microbial pathways to neuronal circuit dynamics, mitochondrial homeostasis and healthspan outcomes with single-organism resolution in a high-throughput manner. For example, commensal *Providencia* bacteria colonizing the worm gut produce the neuromodulator tyramine, which can substitute for host-derived tyramine and is converted to octopamine that acts via the OCTR-1 receptor on ASH nociceptive neurons to modulate aversive olfactory behavior ^18^. This work reveals a gut bacterium-derived neurotransmitter that mimics a host signal to override a sensory decision in a way that benefits both host and microbe. Together, these features position *C. elegans* as an ideal system to mechanistically define how gut microbes and their metabolites reprogram nervous system function and aging trajectories in a whole animal. While much work has been dedicated to identifying bacterial molecules that regulate behavior through canonical neurotransmitter mechanisms, little is known about other microbial metabolites tuning host behavior.

Here, our work identifies mitochondria as a signaling and metabolic hub at the core of microbe–host interactions along the gut–brain axis. We performed a comprehensive high-throughput microbe–*C. elegans* behavioral screen to map bacterial factors that alter organismal behavioral phenotypes. This approach revealed that deletion of specific oxidative phosphorylation (OXPHOS) and iron homeostasis genes in bacteria markedly influenced host behavioral outputs such as movement speed, acting through regulation of enterobactin biosynthesis. Host transcriptional profiling uncovered a ROS-dependent activation of the mitochondrial unfolded protein response (UPR^mt^) when exposed to enterobactin-producing bacterial strains. Genetic modulation of ROS-regulating enzymes and respiratory chain complexes demonstrated that mitochondrial redox balance is a major driver of the microbially induced neuromuscular, healthspan and lifespan phenotypes. These mitochondrial perturbations also triggered cell non-autonomous signaling through neuropeptides, coupling neuronal metabolism to systemic behavioral and metabolic responses.

## RESULTS

### A high-throughput behavioral screen identifies bacterial determinants of host behavior

To identify bacterial genes that are required in the microbiome for normal behavior, we recorded worms that were freely behaving on lawns of mutant bacteria from the 3,985 single-gene knockout Keio collection (Fig. 1a). To avoid developmental effects, worms were raised on wild-type *E. coli* and transferred to wells seeded with mutant bacteria for 4 hours before imaging using camera arrays ^17^. We quantified worm motion, posture and morphology, both at baseline and in response to blue light (an aversive stimulus) using the automated worm tracker Tierpsy ^19^, and compared worms on mutant *E. coli* to worms on wild-type controls on a feature-by-feature basis using two-sided *t*-tests. We selected 52 of the 100 strains with the lowest p-values in any feature, as well as strains that clustered with them to produce a list of 232 strains that we rescreened with more replicates. A heatmap of the data from the confirmation screen shows the phenotypes induced in worms by mutant *E. coli* strains (Fig. 1b). When functionally related *E. coli* genes are knocked out, the resulting mutant bacterial strains have similar effects on worm behavior, which is reflected in how they cluster (dendrogram, Extended Data Fig. 1a,b; Supplementary Table 1). These functional relationships produce a significant enrichment in KEGG pathways and gene ontology terms within the selected clusters (Fig. 1c; Extended Data Fig. 1c, 2a,b).

**Figure 1.**
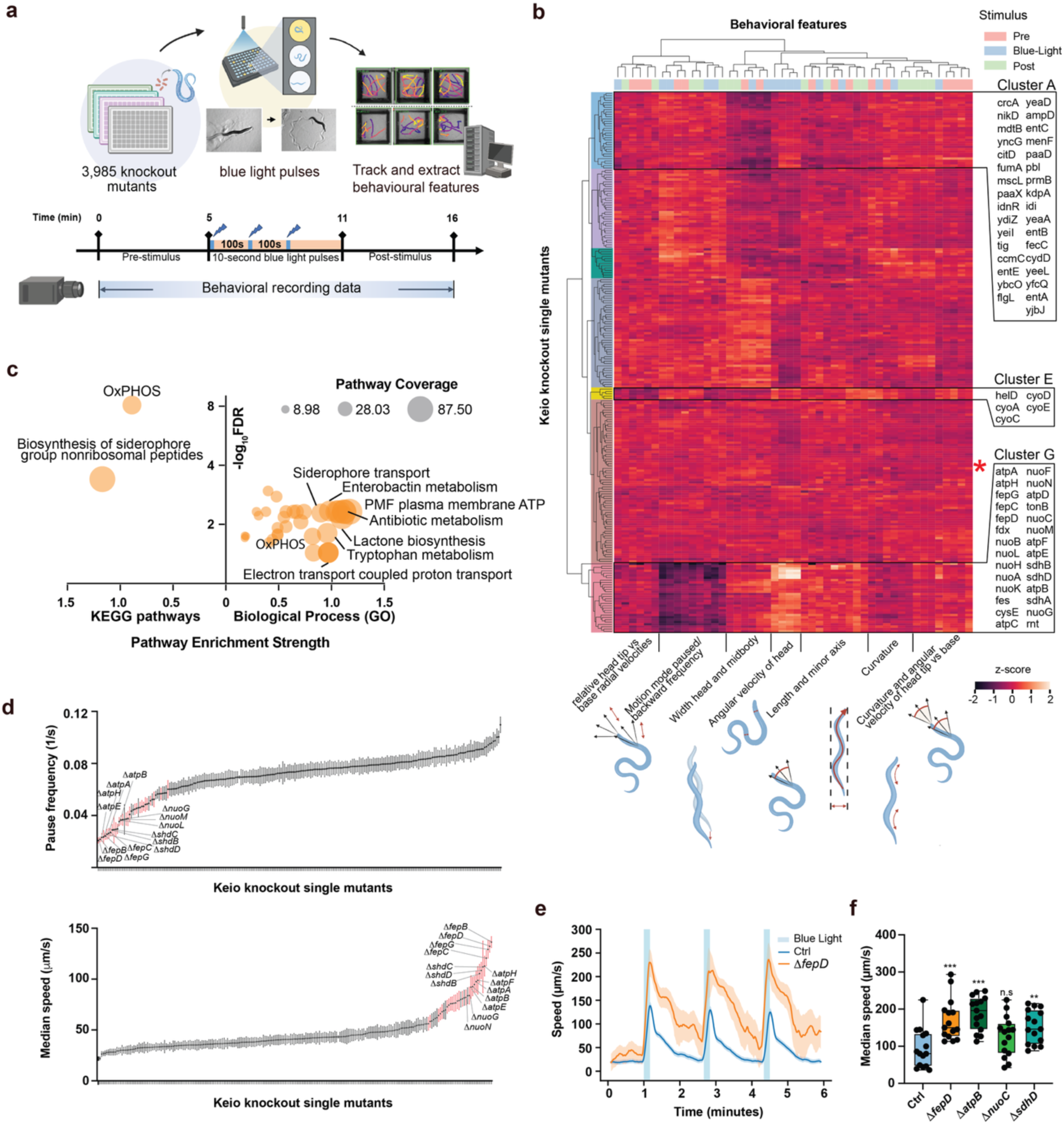
Quantitative tracking reveals clusters of *E. coli* mutations in several pathways that affect host behavior. **a**, Schematic overview of the microbe–host behavioral high-throughput screening (MH-BHTS) platform. *C. elegans* (N2) fed on the *E. coli* Keio single-gene knockout library and the parental strain BW25113. For each bacterial deletion, 3,016 behavioral features were quantified (>12 million data points). **b,** A reduced feature set (Tierpsy16) was used to generate a heatmap of the MH-BHTS dataset, revealing three clusters of bacterial mutants that significantly alter host behavior. The control strain BW25113 is indicated with a red asterisk. **c,** Functional enrichment analysis of mutants exhibiting significant behavioral alterations. **d,** Ranked distribution of pausing frequency under blue-light stimulation, and median speed for each mutant of the Keio library. **e,** Time course of speed in response to blue light stimulation (blue area) with worms fed on control BW25113 (n=174) or Δ*fepD* (n=53) bacteria. **f,** Median speed after blue light stimulation of *C. elegans* fed on control BW25113, Δ*fepD*, Δ*atpB*, Δ*nuoC* or Δ*sdhD* bacteria. Each dot represents the average of worms in one well (n=15 with 10 worms/well). Statistics correspond to comparison of median speed on each mutant bacteria against the control BW25113 (two-sample *t*-tests, FDR controlled at 5% using Benjamini-Hochberg procedure), represented as *p < 0.05, **p < 0.01, ***p < 0.001, n.s p > 0.05.

We focused on cluster G, whose genes had the strongest behavioral effects on worms when knocked out in *E. coli*. Worms exposed to bacteria in this cluster were hyperactive with a marked decrease in pause frequency and an increase in speed upon blue light stimulation. Genes involved in pathways related to the siderophore enterobactin, a small high-affinity iron-chelating metabolite produced by several microbial members belonging to the Enterobacteriaceae family as well as some Actinomycetes strains ^20^, showed particularly strong effects on worms (Fig. 1d). For example, *ΔfepD E. coli*, which have a mutation in the gene coding for the inner membrane permease that imports ferric enterobactin (Extended Data Fig. 2c), led to a higher median speed during blue light videos. By plotting a time series of speed during blue light trials, in addition to a higher speed at baseline, we found worms that were hypersensitive to blue light (Fig. 1e). To specifically quantify this blue light response, we measured worm speed during a 10-s window starting 20 s after the last of three blue light pulses. Speed during this window is used as a phenotype throughout the rest of the study. Bacteria with mutations in enterobactin-related genes, as well as those belonging to subunits of the respiratory chain involved in OXPHOS, led to marked increases in worm speed after blue light stimulation (Fig. 1f). Importantly, our behavioral findings are distinct from those of a development-based screen with the Keio library ^21^ (Extended Data Fig. 2d,e) suggesting novel mechanisms by which bacteria regulate host behavior.

### Bacterial central carbon metabolism regulates enterobactin production controlling worm locomotion speed

To identify bacterial mechanisms regulating host behavior, we performed proteomic profiling of *E. coli* deficient in ferric enterobactin transport (Δ*fepD*) and OXPHOS (Δ*nuoC*, Δ*sdhC* Δ*atpB*). Despite exhibiting largely distinct proteomic signatures with minimal overlap in KEGG functional categories (Extended Data Fig. 3a; Supplementary Table 2), all mutants consistently showed strong upregulation of FecA, encoding an outer membrane receptor for ferric citrate uptake ^18^ (Fig. 2a; Extended Data Fig. 2c). This common response suggests a convergent state of iron limitation in all 4 mutants. To assess the contribution of iron availability, we manipulated iron levels pharmacologically and genetically. Treatment with the iron chelators deferoxamine or citrate decreased worm locomotion speed on both control and Δ*fepD* mutant bacteria (Fig. 2b; Extended Data Fig. 3b). Similarly, deletion of Δ*fecA* decreased worm speed in a Δ*fepD* mutant background, although not fully suppressing the elevated locomotion speed observed with Δ*fepD* mutant bacteria (Fig. 2c). Conversely, iron supplementation (FeSO₄) increased worm speed on control bacteria, but normalized the heightened locomotion speed induced by Δ*fepD* mutants (Fig. 2d). Together, these findings indicate that iron availability alone has a limited direct effect on host behavior, pointing instead to iron-associated bacterial metabolites as key mediators in eliciting this phenotype.

**Figure 2.**
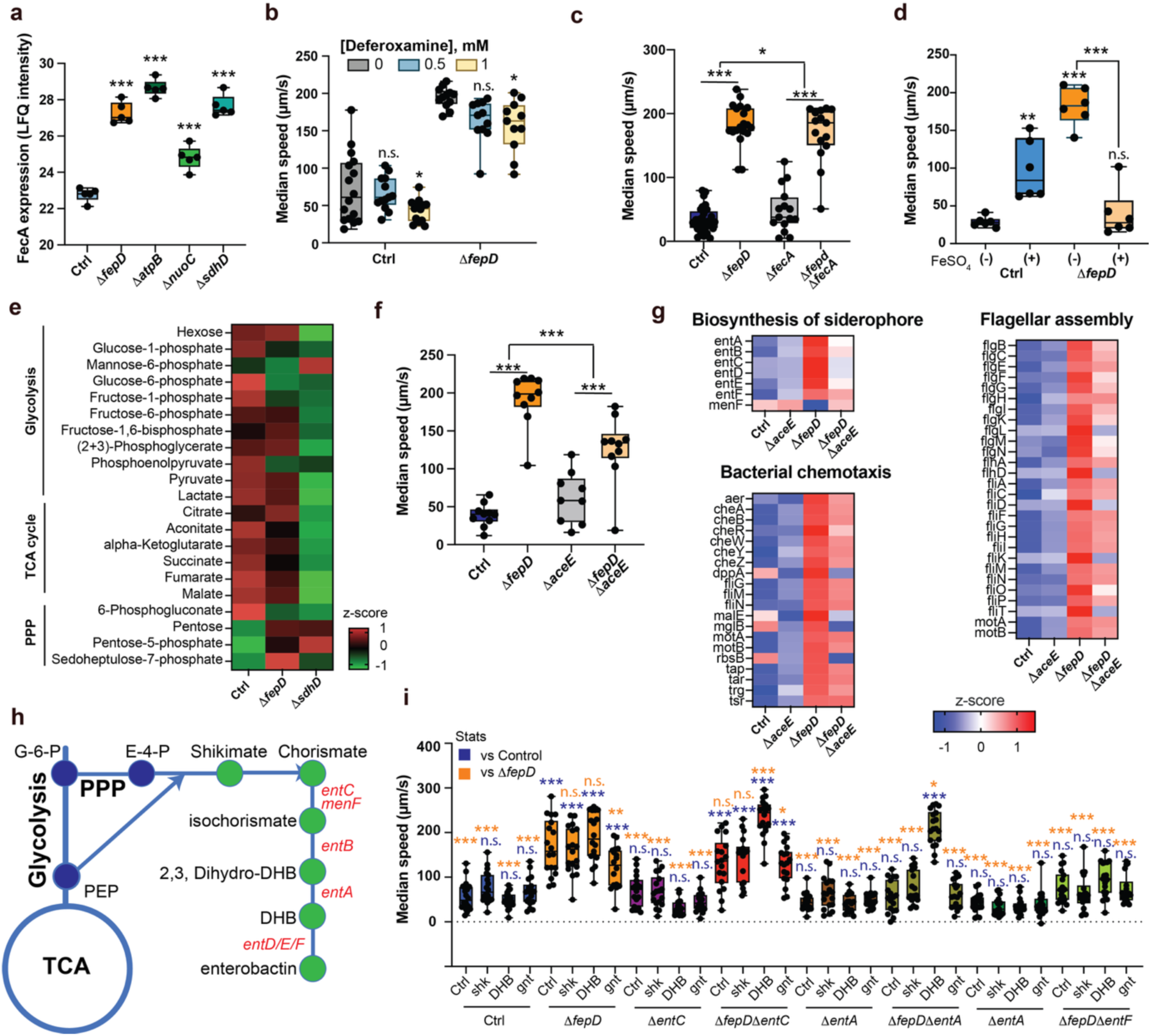
*FepD* bacterial mutants rewire central carbon metabolism for increased enterobactin production and regulation of host behavior. **a,** FecA expression in control BW25113, or Δ*fepD*, Δ*atpB*, Δ*nuoC* and Δ*sdhD* mutants from proteomics data (n=5). **b,** Impact of deferoxamine supplementation (0, 0.5, 1 mM) on *C. elegans* median speed when fed control BW2511 or Δ*fepD* mutant bacteria. The supplementation was compared against the condition without supplementation. Each dot represents the average of worms in one well (n=11-16 with 10 worms/well). **c,** Median speed of *C. elegans* fed on control BW25113, Δ*fepD*, Δ*fecA* or Δ*fepD*Δ*fecA* bacteria. Each dot represents the average of worms in one well (n=15-20 wells with 10 worms/well). **d,** Impact of FeSO_4_ supplementation on worm median speed when fed on control BW25113 and Δ*fepD.* Each dot represents the average of worms in one well (n=6 with 10 worms/well). **e,** Metabolite content from glycolysis, tricarboxylic acid cycle (TCA) cycle, and pentose phosphate pathway (PPP) in control BW25113, Δ*fepD* or Δ*sdhD* bacteria. **f,** Median speed of worms fed on control BW25113, Δ*fepD*, Δ*aceE* or Δ*fepD*Δ*aceE* bacteria. Each dot represents the average of worms in one well (n=9-10 wells with 10 worms/well). **g,** Expression profile of proteins involved in the biosynthesis of siderophore, flagellar assembly and bacterial chemotaxis in control BW25113, Δ*fepD*, Δ*aceE* or Δ*fepD*Δ*aceE* bacteria. **h,** Enterobactin synthesis pathways from glucose-6-phosphate (G-6-P) and shortcut from phosphoenolpyruvate (PEP). **i,** Median speed of worms fed on control BW25113, Δ*fepD*, Δ*entC*, Δ*fepD*Δ*entC*, Δ*entA*, Δ*fepD*Δ*entA*, Δ*entF*, and Δ*fepD*Δ*entF* bacteria, with or without shikimic acid (shk), 2,3-dihydroxybenzoic acid (DHB) or gentisic acid (gnt). Each dot represents the average of worms in one well (n=18 with 10 worms/well). Statistics were performed against the control (one-way ANOVA), between nested conditions (two-way ANOVA), represented as *p < 0.05, **p < 0.01, ***p < 0.001, n.s p > 0.05.

Given the central role of OXPHOS in metabolic flux, we next performed targeted metabolomics. Both Δ*fepD* and Δ*sdhD* mutants displayed marked decreases in glycolytic and TCA cycle intermediates, accompanied by increased pentose phosphate pathway metabolites (Fig. 2e), indicating a substantial rewiring of central carbon metabolism. To functionally link these metabolic changes to host behavior, we leveraged the Δ*fepD* proteomic dataset (Supplementary Table 2) to generate double mutants targeting upregulated metabolic genes and known regulators of iron homeostasis, as well as strains overexpressing downregulated proteins (Extended Data Fig. 3c,d). Among these, only deletion of Δ*aceE,* which encodes the E1 subunit of pyruvate dehydrogenase, which connects glycolysis to the TCA cycle, significantly modulated the Δ*fepD-*induced increase in worm speed (Fig. 2f). Proteomic analysis of control, Δ*fepD*, Δ*aceE*, and Δ*fepD*Δ*aceE* double mutants revealed enrichment of pathways related to siderophore biosynthesis, flagellar assembly, and chemotaxis (Fig. 2g; Extended Data Fig. 4; Supplementary Table 3). Notably, the Δ*fepD*-dependent upregulation of enterobactin biosynthetic enzymes (EntA–EntF) was restored toward baseline in the Δ*fepD*Δ*aceE* double mutant. To directly test the role of enterobactin, we disrupted multiple steps in its biosynthetic pathway and performed metabolite complementation assays (Fig. 2h). Loss of enterobactin production in these double mutants, with and without iron supplemention, was confirmed experimentally (Extended Data Fig. 4d,e). Abolishing enterobactin production fully suppressed the increased locomotion speed induced by Δ*fepD* mutant bacteria (Fig. 2i). Collectively, these results demonstrate that bacterial regulation of host locomotion speed is mediated not by iron availability *per se*, but by the production of iron-chelating siderophores, with enterobactin emerging as a key effector of host behavioral modulation.

### Bacterial enterobactin biosynthesis regulates host UPR^mt^, neuromuscular function and lifespan

To determine how enterobactin influences host physiology, we performed bulk RNA-seq on worms fed control bacteria, enterobactin-producing Δ*fepD* mutants, and non-producing Δ*entF* or Δ*fepD*Δ*entF* strains. The largest transcriptional differences were observed in worms fed Δ*fepD* bacteria (Fig. 3a; Extended Data Fig. 5a,b; Supplementary Table 4). Gene ontology analysis and FDR-ranked biological processes revealed significant enrichment of detoxification pathways, mitochondrial function, muscle integrity, neuronal physiology, and locomotion (Fig. 3b, Extended Data Fig. 5c). Consistent with our analysis, integration with published datasets of predicted *C. elegans* enterobactin interactors ^14^ highlighted muscle-related processes (Extended Data Fig. 5d,e). We validated these findings by examining *myo-3*, a myosin heavy chain expressed in striated muscle. In agreement with our RNA-seq data, Δ*fepD* feeding promoted sustained, age-dependent maintenance of *myo-3* expression, preserved muscle integrity, and improved locomotion during aging (Fig. 3c–e). Given the link between mitochondrial function and aging, we assessed activation of the mitochondrial unfolded protein response (UPR^mt^) using the *Phsp-6*::GFP reporter. Enterobactin overproduction in Δ*fepD* mutants, as well as supplementation with its precursors, shikimate and dihydroxybenzoic acid (DHB), increased UPR^mt^ in a bacterial genotype-dependent manner (Fig. 3f). We next examined lifespan as this phenotype has not previously been reported in the literature, and found that Δ*fepD* robustly extended lifespan. This effect required *entF* (Fig. 3g; Supplementary Table 7) and *entA* (Extended Data Fig. 6a; Supplementary Table 7), but not *entC* (Extended Data Fig. 6b; Supplementary Table 7), consistent with metabolic redundancy between *entC* and *menF* (Fig. 2h; Extended Data Fig. 6c). Similar dependencies were observed for locomotion and UPR^mt^ activation (Extended Data Fig. 6d,e). Lifespan and UPR^mt^ activation by Δ*fepD* were fully dependent on worm *atfs-1*, a stress-activated transcription factor that mediates the UPR^mt^ (Fig. 3h,i; Supplementary Table 7). Enterobactin was also found to activate the integrated stress response in HCT-116 mammalian cells, as shown by the increased phosphorylation of the eukaryotic translation initiation factor 2 alpha (eIF2a) and downstream translation of selected stress-responsive proteins such as ALDH18A1 (Fig. 3j; Extended Data Fig. 6f). In line with these improved functional readouts, Δ*fepD* mutant-fed animals also exhibited decreased age-associated intestinal autofluorescence (lipofuscin), indicating delayed accumulation of age pigment and improved healthspan (Extended Data Fig. 6g). Perturbation of the chorismate super-pathway or precursor supplementation modestly increased UPR^mt^ and lifespan (Extended Data Fig. 6h–j), indicating that upstream metabolites contribute to regulating this pathway but are not sufficient on their own for full phenotypic effects.

**Figure 3.**
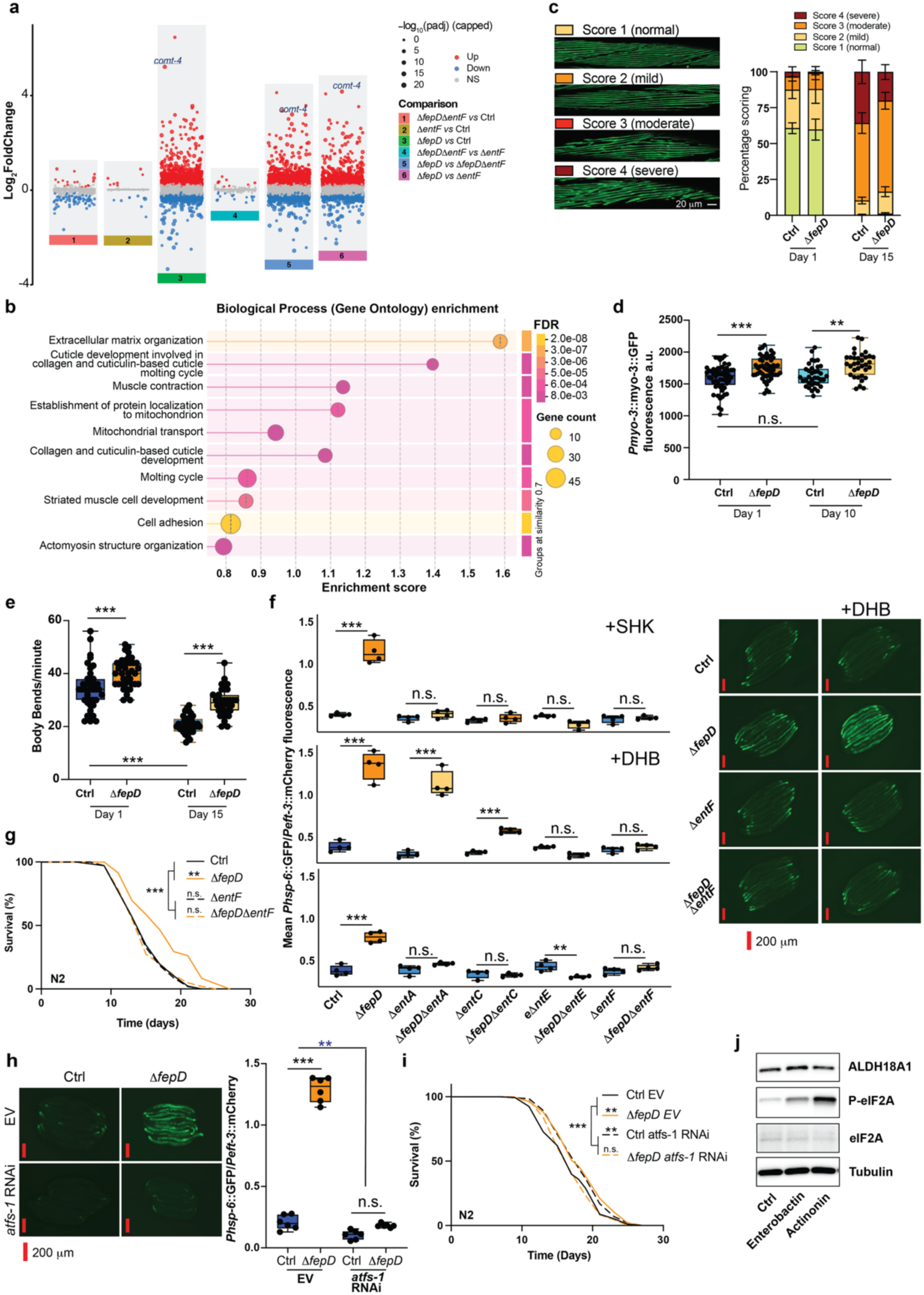
Enterobactin activates UPR^mt^ and improves healthspan and lifespan. **a**, Differential RNA-seq analysis in worms fed on BW25113 control or Δ*fepD*, Δ*entF* and Δ*fepD*Δ*entF* mutants. Dot plots showing gene expression changes across the indicated pairwise comparisons, the dot size is proportional to −log_10_(adjusted P) and each dot represents one gene. Over-expressed and under-expressed genes are represented in red and blue, respectively. Selected genes are labelled. The data were produced from n=5 biological replicates. **b,** Biological process enrichment comparing Δ*fepD* and Δ*fepD*Δ*entF* from RNAseq dataset. **c,** Muscle structure of worms fed on Δ*fepD* and the control BW25113 on day 1 and 15. Worm muscles were blindly scored with n=10 per condition using the worm reporter strain *raIs5 [myo-3p::GFP::myo-3 + rol-6(su1006)]*. **d,** Expression of GFP-tagged Myo-3 protein that localizes to myofilaments in muscle cells of worms fed on the control BW25113 and Δ*fepD* after 1 and 10 days. Each dot represents one worm (n=35-40). **e,** Worm body bend rate on control BW25113 or Δ*fepD* bacteria on day 1 and day 15, using 10 animals per replicate per condition, repeated three times independently. Each dot represents one worm (n=35-40). **f,** Normalized brightness of the worm reporter for *hsp-6* promoter activity as a ratio of GFP over mCherry. Worms fed on control BW25113, Δ*fepD,* Δ*entA,* Δ*fepDentA,* Δ*entC,* Δ*fepDΔentC,* Δ*entE,* Δ*fepD*Δ*entE,* Δ*entF* and Δ*fepD*Δ*entF* bacteria, with or without 2,3-dihydroxybenzoic acid (DHB) and shikimic acid (shk). GFP channeling microscopy images for the control and DHB-supplemented conditions. Each dot represents an average of 15 worms (n=4). **g,** Lifespans of N2 worms fed on control BW25113, Δ*fepD,* Δ*entF* and Δ*entF*Δ*fepD* bacteria (n=119-121, 2 independently run replicates). **h,** Normalized brightness of the worm reporter for *hsp-6* promoter activity as a ratio of GFP over mCherry measured in N2 and *atfs-1* RNAi worms fed on control BW25113 or Δ*fepD* bacteria. Corresponding GFP microscopy images. Each dot represents an average of 15 worms with n=6. **i,** Lifespans of N2 worms fed on *atfs-1* RNAi control BW25113 or the *atfs-1* RNAi Δ*fepD* (n=225-253, 4 independently run replicates). **j,** Protein expression levels upon treatment of HCT-116 cells with 100 μM enterobactin and 100 μM actinonin for 24 h. Microscopy images were analyzed with ImageJ; Each dot represents one worm. Statistics were performed against the control (one-way ANOVA, or Log-Rank for survival curves), between nested conditions (two-way ANOVA or Cox Proportional Hazards for survival curves), represented as *p < 0.05, **p < 0.01, ***p < 0.001, n.s. p > 0.05.

We further investigated *comt-4*, a catechol-O-methyltransferase homolog significantly upregulated in worms fed Δ*fepD* bacteria (Fig. 3a). *comt-4* expression responded to multiple catechol intermediates in the enterobactin biosynthetic pathway, but not to enterobactin itself (Extended Data Fig. 7a,b), and was indirectly influenced by other branches of chorismate metabolism (Extended Data Fig. 7c). However, deletion or knockdown of worm *comt-2, −3, −4, and −5* or *bas-1* (an L-amino acid decarboxylase critical for synthesizing serotonin and dopamine in the nervous system which are detoxified by *comt-4*) did not impair UPR^mt^ activation (Extended Data Fig. 7d) or lifespan extension (Extended Data Fig. 7e,f) by Δ*fepD* mutant bacteria, suggesting that COMT activity, dopamine and serotonin synthesis or breakdown are dispensable for these effects. Instead, *comt-4* may represent a shared detoxification node integrating host-derived catecholamines, such as dopamine, and microbially derived catechols (Extended Data Fig. 7g). Together, these data indicate that metabolites within the enterobactin biosynthetic pathway broadly modulate host physiology, whereas enterobactin itself specifically promotes altered behavior, improved healthspan and lifespan through an as-yet undefined mechanism (Extended Data Fig. 7c,e).

### Enterobactin impacts redox metabolism to regulate host physiology

Next, we investigated the molecular mechanisms through which enterobactin regulates host physiology. We first assessed the role of *atp-1*, the alpha subunit of mitochondrial ATP synthase, previously implicated in mediating mitochondrial import of enterobactin ^14,15^. RNAi knockdown of *atp-1* alone robustly activated UPR^mt^, and exposure to Δ*fepD* mutant bacteria did not further enhance this activation (Extended Data Fig. 8a). In contrast, the effects of enterobactin on *C. elegans* growth were additive to *atp-1* knockdown (Extended Data Fig. 8b), indicating that enterobactin engages additional targets beyond mitochondrial import. To systematically identify candidate protein targets, we performed thermal proteome profiling (TPP), a mass spectrometry-based approach that detects ligand-induced changes in protein thermal stability across the proteome. Enterobactin treatment of worm protein lysates resulted in significant stabilization of 292 proteins and destabilization of 53 proteins, with Δ*T_m_*_50_ shifts ranging from 0.8 to 13.5 and −1.5 to −11.9, respectively (Fig. 4a; Supplementary Table 5). To assess specificity, we conducted a parallel TPP analysis using DHB, a catechol precursor of enterobactin. In contrast to enterobactin, DHB stabilized only 16 proteins and destabilized 286 proteins (Δ*T_m_*_50_: 1.8 to 9.3 and −1.0 to −10.8, respectively; Extended Data Fig. 8b; Supplementary Table S6). Network analysis using STRING revealed largely non-overlapping interaction clusters between the enterobactin and DHB datasets (Extended Data Fig. 8d,e), supporting distinct modes of action despite their structural relatedness. Strikingly, the most strongly stabilized protein identified in the enterobactin dataset was glutaryl-CoA dehydrogenase (GCDH; Δ*T_m_*_50_ = 13.5; Fig. 4b), while two of the most destabilized proteins were the electron-transfer flavoprotein subunits ETFA (Δ*T_m_*_50_ = −11.0; Fig. 4c) and ETFB (Δ*T_m_*_50_ = −11.9; Fig. 4d). These proteins function within the same metabolic reaction, mediating electron transfer from GCDH to the ETF complex (Fig. 4e), suggesting that enterobactin may perturb this redox system. ATP-1 was also stabilized (Δ*T_m_*_50_ = 4.6), consistent with a potential interaction but of lower magnitude.

**Figure 4.**
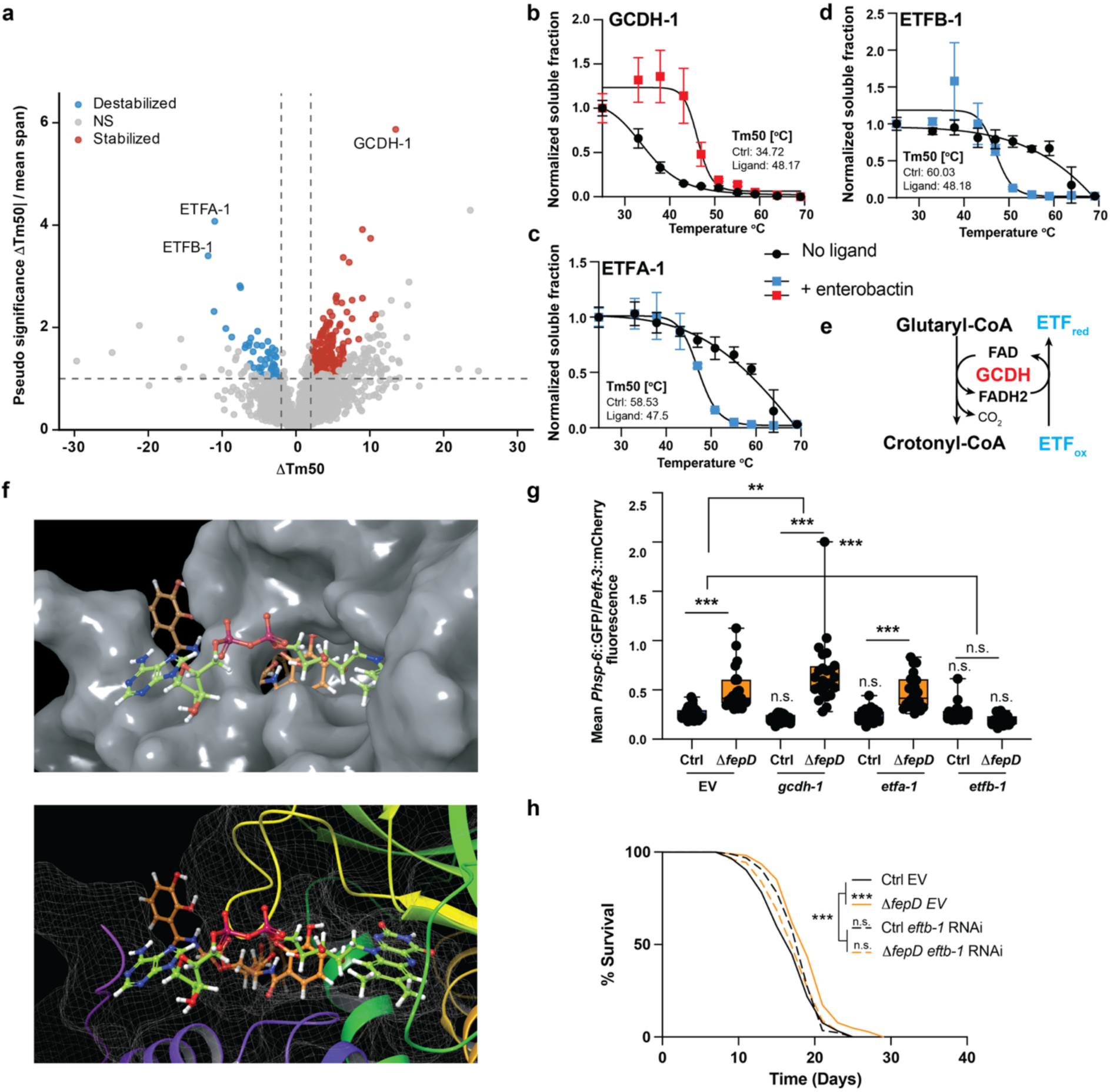
Enterobactin stabilizes mitochondrial glutaryl-CoA dehydrogenase and destabilizes flavoproteins A and B to induce UPR^mt^. **a**, *C. elegans* protein stability changes following treatment with 100 µM enterobactin. Each dot represents one quantified protein. The stability measurement is relative to the melting temperature (Δ*T_m_*_50_) with positive values for protein stabilization and negative values for protein destabilization. These results are expressed as the ratio Δ*T_m_*_50_/mean span. Proteins significantly stabilized are highlighted in red and significantly destabilized proteins in blue. Grey dots denote proteins with insufficient or non-canonical melting curve fits. Dashed lines indicate the thresholds used for significance and effect size. GCDH, glutaryl-CoA dehydrogenase; ETF, electron-transfer flavoprotein. **b-d,** Thermal shift curve for GCDH (**b**), ETFA (**c**) and ETFB (**d**), plotting the normalized soluble fraction against temperature for control and ligand (enterobactin)-treated conditions (n=3). **e,** Reaction catalyzed by GCDH. GCDH converts glutaryl-CoA into crotonyl-CoA, coupled to decarboxylation and reduction of ETF, generating CO_2_. **f,** Embedding of enterobactin (orange) and the FAD cofactor (green) within the structure of GCDH by SwissDock. Top - Surface representation of the GCDH binding pocket with the ligands modelled in stick representation, highlighting its extended conformation within the groove and illustrating potential contact sites along the pocket. Bottom - Ribbon representation of the protein with the ligands shown occupying a long, curved groove. The binding pocket is visualized as a mesh surface to illustrate the enclosing cavity. **g,** Normalized brightness of the worm reporter for *hsp-6* promoter activity as a ratio of GFP over mCherry. Worms were fed on control BW25113 or Δ*fepD* bacteria carrying the control plasmid (EV) or a plasmid for the knockdown of *gcdh-1, etfa-1 or etfb-1.* Each dot represents one worm (n=24, with two independent replicates). **h,** Lifespans of N2 worms fed on *etfb-1* RNAi control BW25113 or *etfb-1* RNAi Δ*fepD* bacteria (n=125-153, 4 independently run replicates). Statistics were performed against the control (one-way ANOVA, or Log-Rank for survival curves), between nested conditions (two-way ANOVA or Cox Proportional Hazards for survival curves), represented as *p < 0.05, **p < 0.01, ***p < 0.001, n.s p > 0.05.

To explore the structural basis of this interaction, we performed molecular docking using SwissDock (Attracting Cavities 2.0 protocol)^22^. The top-ranked binding pose positioned enterobactin adjacent to the FAD cofactor within GCDH (Fig. 4f), suggesting that enterobactin could interfere with electron transfer at the level of flavin-dependent catalysis. To functionally validate these findings, we depleted components of this pathway by RNAi. Knockdown of *etfB-1* completely abolished the UPR^mt^ induction triggered by Δ*fepD* bacteria, whereas knockdown of *gcdh*-*1* further enhanced this response (Fig. 4g). Similar genetic interactions were observed for organismal growth and lifespan (Fig. 4h; Extended Data Fig. 8h; Supplementary Table 7), supporting a model in which enterobactin modulates host physiology by disrupting electron transfer between GCDH and the ETF complex.

Since flavoproteins ETFA/B transfer electrons to the respiratory chain, we tested whether knocking out diverse complexes of the electron transport chain (ETC) regulates host locomotion speed. Functional Complexes I–III of the respiratory chain were required for the effect of *ΔfepD* mutants on host locomotion speed, indicating that an intact ETC is necessary for this phenotype (Fig. 5a). Complex I impairment by *gas-1* mutation similarly abolished both Δ*fepD*-mediated lifespan extension and further UPR^mt^ activation (Extended Data Fig. 9a,b). Mitochondrial electron-transfer flavoproteins ETFA/B have been shown *in vitro* to generate substantial superoxide and hydrogen peroxide in conjunction with medium-chain acyl-CoA dehydrogenase (MCAD), with disease-associated ETF variants and altered protein–protein interactions enhancing this ROS production ^23^. To test whether enterobactin-dependent modulation of electron transfer engages a similar mechanism *in vivo*, we combined pharmacological and genetic perturbations of redox homeostasis. Treatment with the ROS scavengers N-acetyl cysteine (NAC) and vitamin C fully suppressed the Δ*fepD*-dependent increase in locomotion speed (Fig 5b, Extended Data Fig. 9c). Likewise, overexpression of the mitochondrial superoxide dismutase *sod-2*, which in *C. elegans* is predominantly expressed in neurons ^24^, but not other antioxidant enzymes, completely abolished the effects of Δ*fepD* on locomotion (Fig. 5c), decreased UPR^mt^ activation (Fig. 5d), and lifespan extension (Fig. 5e), whereas *sod-2* loss-of-function produced the opposite phenotypes (Fig. 5d,f). Consistent with a role for enterobactin in promoting ROS production, worms carrying the rxRFP redox-sensitive RFP reporter displayed increased fluorescence when fed Δ*fepD* mutant bacteria (Fig. 5g), indicative of a more oxidizing intracellular environment. Similarly, supplementation with paraquat, which can accumulate in the mitochondrial matrix and undergoes redox cycling at Complex I to drive superoxide formation ^25^, phenocopied the enterobactin-dependent effects on locomotion in a host-genotype-dependent manner that required the respiratory chain and *sod-2* (Extended Data Fig. 9d-h). Manipulation of host redox capacity by overexpression of catalases CTL-1, CTL-2, and CTL-3 in *C. elegans*, doubled the lifespan of worms fed Δ*fepD* mutant bacteria, an effect that was not additive with *sod-2* deletion (Fig 5h,i; Supplementary Table 7), implying epistatic functional overlap.

**Figure 5.**
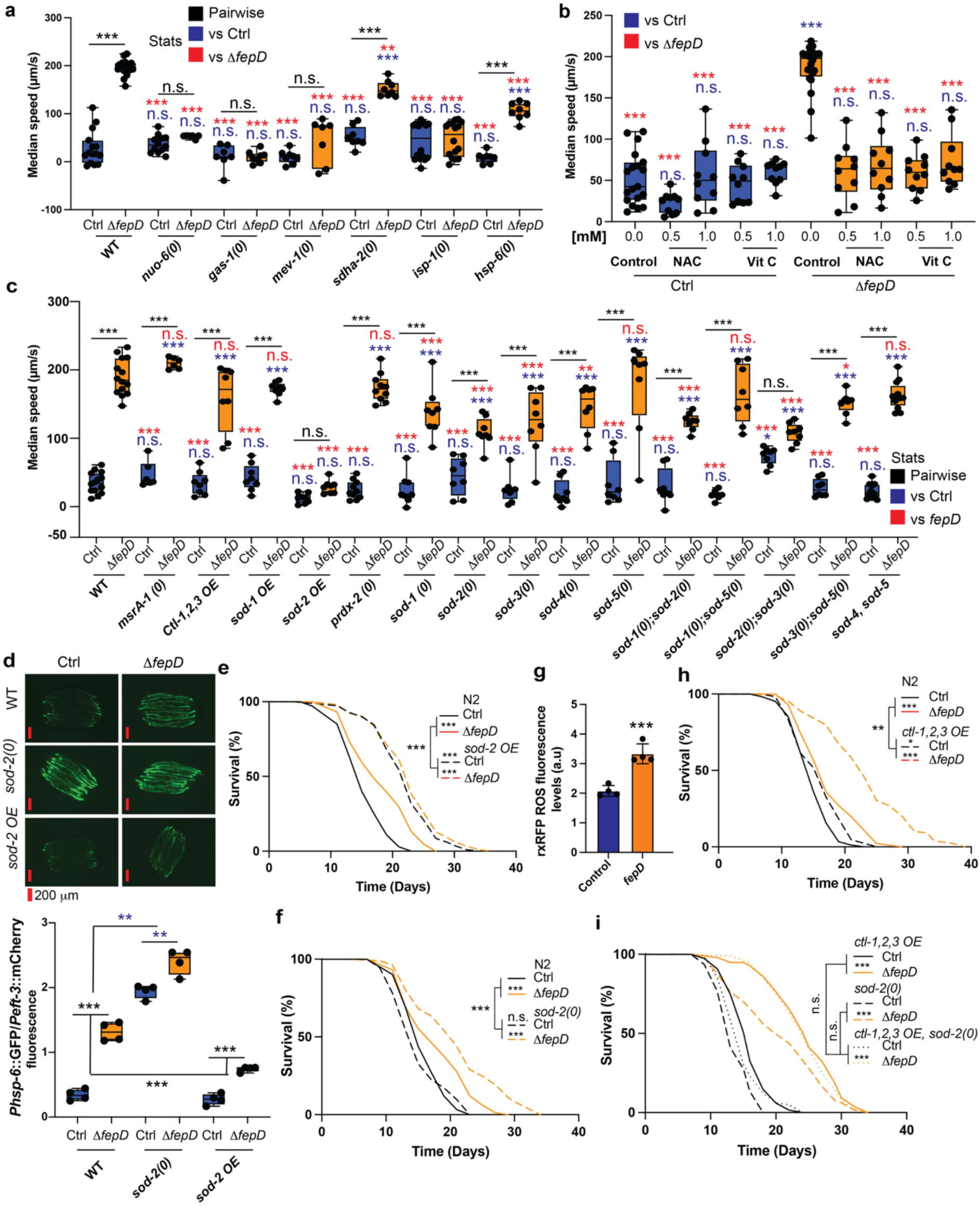
Redox-dependent induction of UPR^mt^, lifespan and behavior. **a**, Median speed measured in wild-type N2 and mitochondrial null mutant *C. elegans* (*nuo-6, gas-1, mev-1, sdha-2, isp-1, hsp-6*) fed on control *E. coli* BW25113 or Δ*fepD* bacteria. Each dot represents the average of worms in one well (n=6-16 wells with 10 worms/well). **b,** Supplementation of vitamin C and N-acetylcysteine (NAC) (0.5 or 1 mM) on worms fed control BW25113 or Δ*fepD* mutant bacteria. Each dot represents the average of worms in one well (n=10-20 wells with 10 worms/well). **c,** Speed measured in N2 and oxidative stress-related mutant *C. elegans* (deletion of *msra-1, ctl-1/2/3, prdx-2, sod-1, sod-2, sod-3, sod-4, sod-5,* overexpression (OE) of *sod-1* and *sod-2*, *ctl-1/2/3* and indicated combinations) fed on control BW25113 or Δ*fepD* bacteria. Each dot represents the average of worms in one well (n=6-14 wells with 10 worms/well). **d,** Normalized brightness of the worm reporter for *hsp-6* expression as a ratio of GFP over mCherry measured in N2, *sod-2* null mutant, and *sod-2* overexpressing worms fed on control BW25113 or Δ*fepD* bacteria. Each dot represents an average of 15 worms with n=4. **e,f,** Lifespans of N2 worms and worms with *sod-2* overexpression (n=120-125, 2 independently run replicates) (**e**) and *sod-2* null mutation (n=118-124, 2 independently run replicates) (**f**) fed on control BW25113 or Δ*fepD* bacteria. **g**, ROS fluorescence level for N2 worms fed on control BW25113 or Δ*fepD* bacteria. Each dot represents an average of 15 worms with n=4. **h, i,** Lifespans of N2 and *ctl-1,2,3* overexpressing worms (n=119-126, 2 independently run replicates) (**h**), and comparison between worms with *sod-2* null mutation alone, or combined with *ctl-1,2,3* overexpression (n=112-151, 2 independently run replicates) (**i**) fed on control BW25113 or Δ*fepD* bacteria. Microscopy images were analyzed with ImageJ. Statistics were performed against the control (one-way ANOVA, or Log-Rank for survival curves), between nested conditions (two-way ANOVA or Cox Proportional Hazards for survival curves), represented as *p < 0.05, **p < 0.01, ***p < 0.001, n.s p > 0.05.

Together, these data suggest that enterobactin-dependent modulation of GCDH-ETFB flavoprotein electron transfer to the respiratory chain induces redox changes in neuronal mitochondria to tune UPR^mt^ activity, locomotor behavior, and lifespan.

### A secretin-family GPCR controls host behaviour and lifespan in response to enterobactin

In mammals, 3-hydroxyglutarate (3-OHGA) and glutarate, which accumulate upon GCDH dysfunction, can behave as “false neurotransmitters” at glutamatergic synapses, acting as weak N-methyl-D-aspartate (NMDA) receptor agonists and potentially driving NMDA-dependent neuronal overactivation ^26,27^. To test whether enterobactin acts through a similar mechanism, we used MK-801 (dizocilpine), a non-competitive NMDA receptor antagonist that blocks open NMDA receptor channels ^28^. MK-801 not only failed to rescue the effects of Δ*fepD* bacterial mutants on either UPR^mt^ activation, growth impairment and lifespan in *C. elegans*, but rather enhanced all three phenotypes instead (Extended Data Fig. 10a,b,c; Supplementary Table 7), suggesting that enterobactin does not primarily signal via NMDA-type glutamatergic transmission. Thus, to identify neuronal pathways mediating enterobactin effects, we assayed locomotion speed in ASJ and AIY neuron–ablated animals and mutants affecting locomotory, arousal and lethargus-related behaviors. Loss of the Phe-Met-Arg-Phe-NH_2_ (FMRFamide)-like neuropeptide *flp-2* and of the pigment-dispersing factor receptor *pdfr-1* abolished the effects of both Δ*fepD* and paraquat on locomotion speed (Fig. 6a; Extended Data Fig. 10d). Deletion of *pdfr-1* markedly reduced the impact of Δ*fepD-*feeding on growth (Fig. 6b), development (Extended Data Fig. 10e) and UPR^mt^ activation (Fig. 6c), fully suppressed changes in muscle fiber *myo-3* expression (Fig. 6d) and associated healthspan benefits (Fig. 6e). *pdfr-1* knockout also abolished lifespan extension induced by either Δ*fepD* (Fig. 6f; Table S7) or Δ*fepD*Δ*entC* (Extended Data Fig. 10f, Table S7) bacterial mutants, as confirmed using two independent *pdrf-1* alleles (Extended Data Fig. 10g,h, Table S7), and complementation of *pdfr-1* in all neurons or specifically in *tdc-1*, *glr-3* and *ttx-3*-positive neurons restored these effects (Extended Data Fig. 10i, Supplementary Table 7). FLP-2 acts in concert with the pigment-dispersing factor (PDF-1) neuropeptide system to promote arousal and increase locomotor activity ^29^. Consistent with this, we found that *pdf-1* was strictly required for Δ*fepD*-dependent longevity (Fig. 6g; Supplementary Table 7), whereas *flp-2* was dispensable for lifespan regulation (Extended Data Fig. 10j; Supplementary Table 7), indicating a functional divergence between these two arousal-related peptides in this context. Finally, we asked whether *pdfr-1* was required in redox-sensitized genetic backgrounds. Deletion of *pdfr-1* strongly suppressed the long lifespans of *sod-2* null mutants (Extended Data Fig. 10k; Supplementary Table 7) and of *ctl-1,2,3* overexpressor strains (Fig. 6h; Supplementary Table 7) when worms were fed Δ*fepD* mutant bacteria. Overall, these data indicate that enterobactin does not act via canonical NMDA receptor-mediated glutamatergic signaling or catechol-mediated neurotransmitter signaling, but instead engages a *pdf-1/pdfr-1-*dependent peptidergic circuit to couple bacterial redox state to neuronal arousal pathways that coordinately regulate locomotion, growth, UPR^mt^ activation, healthspan and longevity in *C. elegans*.

**Figure 6.**
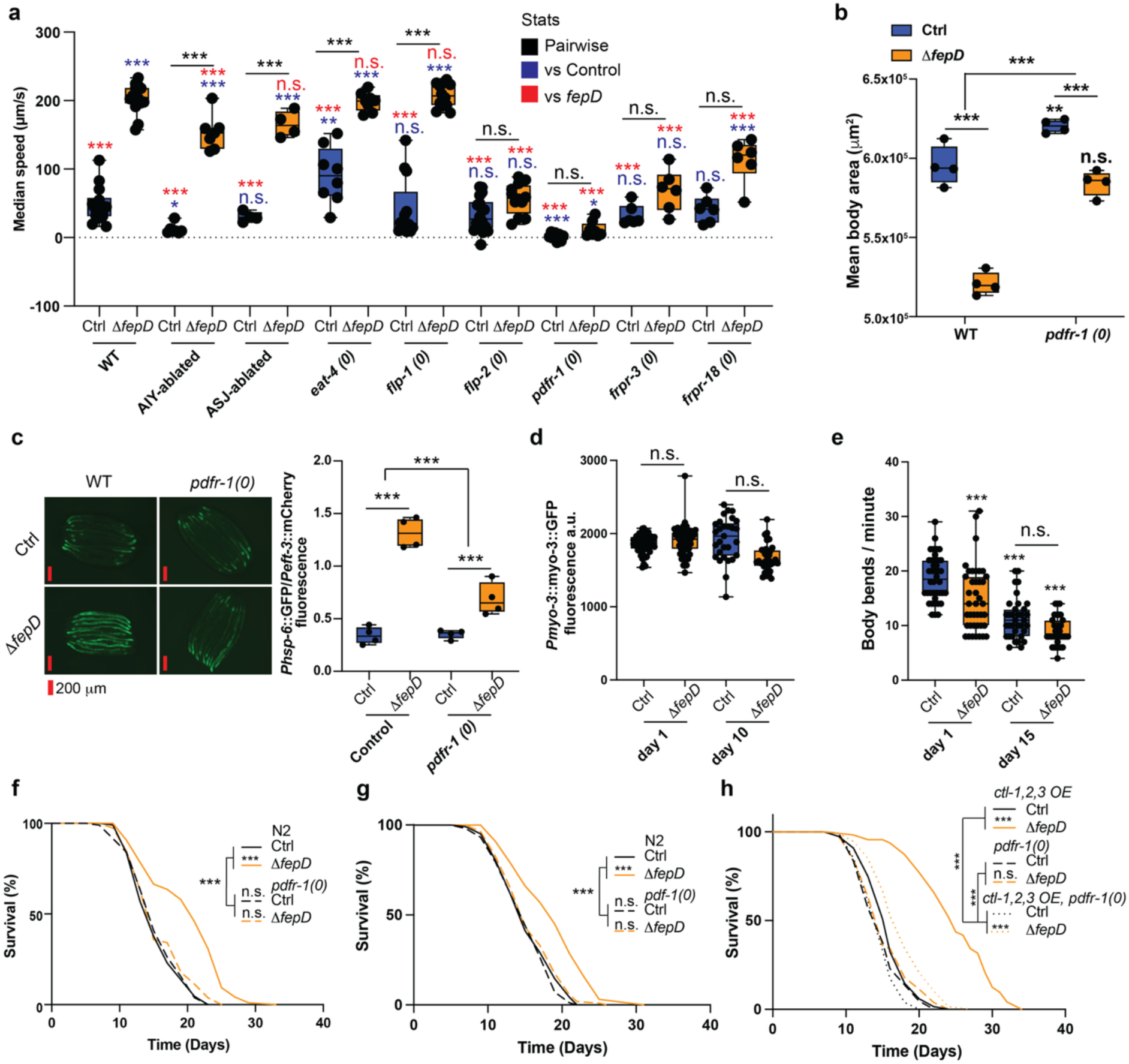
Neuropeptide signaling by PDFR-1 mediates behavioral and lifespan effects induced by enterobactin. **a**, Median speed measured in *C. elegans* N2 wild-type (WT) and neuropeptidergic pathway mutants affecting specific neurons (ASJ, AIY) and signaling components (*eat-4*, *flp-1, flp-2, frpr-3, frpr-18, pdfr-1*) fed on control BW25113 or Δ*fepD* mutant bacteria. Each dot represents the average of worms in one well (n=4-18 wells with 10 worms/well). **b,** Mean body area of N2 and *pdfr-1* null mutant worms fed on control BW25113 or Δ*fepD* bacteria from the L1 stage to adulthood. Each dot represents an average of 15 worms with n=4. **c,** Normalized brightness of the worm reporter for *hsp-6* expression as a ratio of GFP over mCherry measured in N2, *pdfr-1*-null mutant fed on control BW25113 or Δ*fepD* bacteria. Each dot represents an average of 15 worms with n=4. **d,** Expression of GFP-tagged *myo-3* protein that localizes to myofilaments in muscle cells of *pdfr-1-*null mutants worms fed on control BW25113 or Δ*fepD* bacteria after 1 and 10 days. Each dot represents one worm with n=29-55. **e,** Body bends assay of *pdfr-1-*null mutants fed on control BW25113 or Δ*fepD* bacteria, scored for 1 min at day 1 and 15. Each dot represents one worm with n=35-40. **f-h,** Lifespans of N2 and *pdfr-1-*null mutant *C. elegans* (n=118-128, 2 independently run replicates) (f), N2 and *pdf-1* null mutant *C. elegans* (n=119-123, 2 independently run replicates) (g), and *pdfr-1* null mutant and *ctl-1,2,3* overexpression mutant *C. elegans* and their combination (n=127-157, 2 independently run replicates) (h), fed on control BW25113 or Δ*fepD* bacteria. Microscopy images were analyzed with ImageJ. Statistics were performed against the control (one-way ANOVA, or Log-Rank for survival curves), between nested conditions (two-way ANOVA or Cox Proportional Hazards for survival curves), represented as *p < 0.05, **p < 0.01, ***p < 0.001, n.s. p > 0.05.

## DISCUSSION

A central challenge in understanding microbial effects on physiology has been the trade-off between experimental throughput and the richness of the host phenotypes that can be measured. The Keio library of *E. coli* knockouts has been screened in worms for effects on development ^21^, social ^30^ and aversive behaviors ^31^, and lifespan ^32,33^, but these phenotypes were quantified using a single parameter which provides limited information on the functional relationships between hits. The advance of megapixel camera arrays has enabled comprehensive screening of the Keio library with multiple behavioral endpoints simultaneously. Clustering based on these behavioral fingerprints revealed functionally related groups of bacterial genes, involved in the production and transport of two structurally analogous iron-binding metabolites, protoheme and enterobactin. Protoheme is required for cellular respiration in muscle and nerve cells, with severely heme-deprived worms displaying sluggish, paralyzed or arrested phenotypes ^34^. Our behavioral data are consistent with these observations, where worms growing on bacterial mutants involved in protoheme utilization and export (Clusters E and F, Fig 1b) displaying similar behavioral phenotypes. In parallel, the bacterial siderophore enterobactin can deliver iron to *C. elegans* mitochondria supporting normal development ^14,21^. Building on this, our data uncover a central role for gut–brain axis mechanisms in regulating previously unappreciated iron-dependent phenotypes, including specific behavioral states and lifespan. The behavioral specificity we observe suggests that *C. elegans* can discriminate between chemically related iron chelators of bacterial origin, enabling the microbiota to selectively tune discrete neural and behavioral states through subtle differences in iron-binding chemistry rather than iron supply *per se*.

Together, our findings, in conjunction with clinical observations from patients with glutaric acidemia type I (GA-I), a metabolic disorder caused by glutaryl-CoA dehydrogenase (GCDH) deficiency, point to a conserved vulnerability within mitochondrial electron transfer circuitry centered on the GCDH–ETF axis. In humans, pathogenic variants in GCDH impair enzymatic activity, leading to disrupted mitochondrial metabolism promoting glutamatergic excitotoxicity ^35^, ultimately driving the emergence of dystonic–dyskinetic movement disorders ^36^. Our data suggest that this axis is not only genetically susceptible, but also environmentally vulnerable. In *C. elegans*, enterobactin exposure phenocopies behavioral alterations, including increased locomotion and heightened arousal, by selectively stabilizing GCDH and destabilizing ETF subunits. This asymmetric effect on the dehydrogenase–electron acceptor pair is consistent with impaired electron flux into the respiratory chain, and is further supported by the induction of an *etfb-1*-dependent UPR^mt^. ETF complexes generate anion superoxide and hydrogen peroxide when electrons leak from their FAD cofactor to oxygen during transfer, a process amplified by metabolic overload of the respiratory chain and by structural instability or misfolding that increases flavin reactivity ^23^. Consistent with this model, we find that neuronal overexpression of superoxide dismutase 2 (SOD-2) or antioxidant supplementation is sufficient to suppress enterobactin-induced phenotypes, including peripheral UPR^mt^ activation, increased locomotor activity, and lifespan extension. Conversely, genetic manipulations that cause an oxidising shift in redox state, such as *sod-2* deletion ^37^, catalase overexpression ^38^, or supplementation with MK-801 which induces neuronal ROS ^39^, can potentiate enterobactin sensitivity, while paraquat supplementation phenocopies the effects of enterobactin, reinforcing the conclusion that redox imbalance is a key driver of the observed physiological responses to enterobactin. Together with prior work showing that neuronal mitochondrial ROS can drive peripheral UPR^mt^ and regulate digestive physiology ^40^, and that mitochondrial hydrogen peroxide promotes neuropeptide secretion ^41^, our data support a model in which neuronal mitochondrial redox signaling acts as an intermediate of cellular and organismal regulation, rather than a simple by-product of dysfunction.

Within this framework, we identify the neuropeptide ligand–receptor pair PDF-1/PDFR-1 as a critical downstream effector of enterobactin-induced mitochondrial redox signaling. PDF-1, a neuropeptide expressed in a specific set of sensory and interneurons, and its broadly expressed receptor, the GPCR PDFR-1, which mediates cell-non-autonomous signaling, are known regulators of arousal, locomotion, and behavioral state transitions ^42,43^. We propose that redox changes generated upon disruption of the GCDH–ETF axis by enterobactin regulate the PDF-1/PDFR-1-controlled neuromodulatory network that sustains forward motor drive and arousal. Activation of PDFR-1 signaling in peripheral tissues then induces UPR^mt^, linking neuronal redox signaling to organism-wide adaptation. This establishes a mechanistic route by which microbial manipulation of mitochondrial electron transfer is converted into endocrine-like control of behavior and longevity.

Our data collectively argue that gut microbes can act as active architects of host locomotion, with consequences for later-life health. SCFAs exemplify how classical fermentation products are repurposed as signals, linking microbial metabolism of dietary fiber to neuro–immuno–endocrine circuits that regulate appetite and energy balance ^44^. Another growing set of studies shows that specific commensal bacteria synthesize *bona fide* neurotransmitters or neuropeptide mimetics, which interface directly with host canonical neurotransmitter circuits controlling food choice, affect behavior, and locomotion. In this light, our finding that the siderophore enterobactin produced by *E. coli* hijacks the *pdf-1/pdfr-1* neuropeptide axis, known to be involved in foraging and exploratory behavior ^43,45^, to tune neuromuscular tone and aging adds a novel distinct mechanistic layer. Here, a bacterial iron-scavenging system is wired into host motor programs, plausibly biasing animals toward iron-rich niches in which the bacterium itself can better compete. Since enterobactin synthesis is tightly induced by iron limitation, and iron is both essential and potentially toxic ^46^, this neuromodulation may reflect a finely balanced microbial strategy for survival and dispersion ^42^. By enhancing mobility and altering exploration when iron is scarce, bacteria could increase the likelihood that their host encounters microenvironments with more favorable metal availability, while the host gains improvements in neuromuscular performance and healthspan as an unexpected but positive by-product. Conceptually, enterobactin is a novel member of a broader class of “neuro-manipulators” produced by bacteria, distinct from canonical neurotransmitters such tyramine ^47^, and serotonin-inducing consortia ^48^, or α-melanocyte-stimulating hormone-mimetic proteins such as ClpB ^49^, that align or misalign microbial fitness with host behavioral and physiological outputs. We propose that such metabolites and peptide mimetics constitute an underappreciated axis of host–microbe interaction in which evolutionary interests unexpectedly converge or diverge, with microbial signals co-opting conserved neuromodulatory hubs to steer exploration, feeding, stress responses, and now aging trajectories.

## ACKNOWLEDGEMENTS

FC was supported by the Wellcome Trust/Royal Society (102532/Z/12/Z and 102531/Z/13/A), the DFG German Research Foundation (EXC 2030 - 390661388), (IPFP-B02 Filipe Cabreiro) of the Center for Molecular Medicine Cologne and i-HEAD“ (PB-NRW-2022-ANTRAG-025). This work was supported by the Medical Research Council through grant MC-A658-5TY30 to AEXB and MC-M302-M3450 to HMC. Research in the lab of TMH is funded by the DFG under Germany’s Excellence Strategy (EXC 2048/1 - 390686111). The proteomics unit in TMH’s lab (timsTOF-HT, Bruker Daltonic) is funded via DFG-INST 216/1290-1 FUGG. We acknowledge computational resources and support provided by the Imperial College Research Computing Service (http://doi.org/10.14469/hpc/2232). We thank the CECAD Proteomics Facility for proteomics analyses and Christina Mack for support with TPP. We thank Arantza Barrios for sharing the PDFR-1 CRE-LOX strains. We thank Marijana Croon and Chao Wu for advice on ISR western blotting.

## AUTHOR CONTRIBUTIONS

YW, SM and CB have contributed equally. FC and AB conceptualized the research with contributions by YW, SM and CB. CA, DWD, BH, IK, AM and TH performed TPP and proteomics identification and analysis. SB performed metabolomic identification and analysis. YW, SM, CB, AK, AZ, RB, TVP and AA contributed to experimental procedures. JS, A-MW and SK-S performed docking analysis. YW, AB and FC wrote the manuscript with contributions from all authors. YW, CB, HMC, AB and FC participated in the interpretation of the main findings. AB and FC supervised the research and secured funding. All authors read and approved the final manuscript.

## MATERIAL AND METHODS

### *C. elegans*, bacterial strains, and mammalian cell lines used in this study

The nematode and bacterial strains used in this study are listed in the Key Resources Table. *C. elegans* strains generated as part of this study were obtained through genetic crossing and confirmed by PCR. Bacterial mutants were confirmed by PCR. A list of primers used can also be found in the Resources Table. The *E. coli* Keio single-gene deletion library ^50^ (odd numbered strains) and the ASKA Overexpression Collection ^51^ were obtained from the National BioResource Project (NBRP). The RNAi library of *C. elegans* cloned into the feeding vector pL4440 was obtained from Ahringer ^52^ and Vidal libraries ^53^.

HCT116 human colorectal carcinoma cells (ATCC-CCL-247) were cultured in RPMI 1640 medium (Gibco) supplemented with 10% (v/v) fetal bovine serum (FBS; Sigma-Aldrich) and 1% (v/v) penicillin–streptomycin (Gibco). Cells were maintained at 37°C in a humidified incubator with 5% CO_2_. All experiments were performed using cells between passages 20 and 30.

### Nematode growth conditions

*C. elegans* strains were cultured on Nematode Growth Medium (NGM) plates ^54^ at 20°C, fed with 3-day-old bacterial lawns. Worms were synchronized by standard hypochlorite bleaching (4 M NaOH: bleach = 8:7) and allowed to hatch overnight in M9 buffer to obtain age-matched L1 larvae prior to experiments. For strain generation, male worms were generated by heat shock, in which 30 L4-stage hermaphrodites were incubated at 30.5°C for 5–6 h, followed by transfer of 10 worms per bacterial seeded NGM plate each day for 3 consecutive days. Male progeny was identified among the offspring and maintained by repeated crossing with L4-stage hermaphrodites. Males were then crossed with target *C. elegans* strains carrying the desired genetic modifications. Successful crosses were identified by fluorescence markers and/or PCR genotyping. For RNAi feeding experiments, worms were transferred onto 1-day-old bacterial lawns grown on NGM agar supplemented with ampicillin (100 µg/mL; Fisher Scientific) and IPTG (final concentration 1 mM; Thermo Scientific).

### High-throughput behavior screening

#### Bacteria and imaging plate preparation

Bacterial strains were transferred from stock plates into Nunc 96-well polystyrene round-bottom microwell plates containing LB supplemented with 50 mg/L kanamycin and incubated overnight at 37°C, then a small volume of the cultures was transferred using a 96-pin replicator into 200 µL LB without kanamycin before a second round of overnight culture at 37°C to avoid adding kanamycin to the test plate with *C. elegans*. For behavior recordings, 96-square-well plates containing NGM agar media were seeded with 10 µL of liquid culture with an Integra VIAFLO 96/384 (INTEGRA Biosciences Ltd, UK). The plates were wrapped in wet tissue paper to avoid over-drying the agar and incubated for 8 h at 25°C.

N2 *C. elegans* were raised on NGM agar plates seeded with BW25113 wild-type *E. coli* until the first day of adulthood. Worms were collected with PBS 1X and washed 3 times. 3 worms per well were dispensed into the seeded 96-well imaging plates using a COPAS 500 Flow Pilot worm sorter (Union Biometrica). The imaging plates containing worms were then incubated for 4 h at 20°C prior to imaging. The initial screen of the Keio library was replicated 6 times, with each replicate conducted on a separate day.

#### Behavioral phenotyping of C. elegans

Imaging was performed using custom multi-camera imaging rigs that record simultaneously from all wells of 96 well plates ^17^. Each experiment consisted of three videos: a 5 min pre-stimulus video, followed by a 6 min blue light video (with three blue light stimuli delivered in 10-s bursts approximately 100 s apart), followed by a 5 min post-stimulus video. Videos were recorded at 25 frames per s.

#### Analysis of video recordings

Video recordings were analyzed using the Tierpsy Tracker software (version 1.5.3) ^19,55^ available at https://github.com/Tierpsy/tierpsy-tracker. Before calculating the Tierpsy256 behavioral features ^55^, we only kept skeletonized objects with lengths between 700-1300 µm and widths between 20-200 µm that were tracked for more than 1 s and travelled more than 124 µm throughout the video. After calculating feature summaries for each well, we dropped wells where fewer than 80% of features could be calculated or where fewer than 6000 worm skeletons were present across all three videos (pre-stimulus, blue light, and post-stimulus). Further, we removed features if there were more than 5% missing values for that feature across all samples or if there was zero standard deviation for that feature across all samples. Any remaining missing values were imputed with the global mean value for that feature across all samples.^51^ Statistical analyses were performed in Python (version 3.6) using the following packages: *scipy* (version 1.9.0), *sklearn* (version 1.1.2), *numpy* (version 1.23), *pandas* (version 1.4.3), *pathlib* (version 3.10.6), *matplotlib* (version 3.5.2), and *seaborn* (version 0.11.2).

#### Selection of candidate strains for rescreening

To select candidate mutants for confirmation screening, we did feature-by-feature *t*-tests comparing experimental strains to the *E. coli* BW25113 wild-type control. From the 100 strains with the lowest p-value of any feature, we manually selected 52 strains for follow-up. For each of these 52 strains, we found their three nearest neighbors based on the Euclidean distance between their phenotypic fingerprints. Finally, a handful of strains with mutations in pathways related to the hit strains were manually added. This produced a list of 232 strains that were re-screened.

#### Hierarchical clustering analysis for Gene Ontology (GO) enrichment analysis

Hierarchical clustering (Euclidean distance, complete linkage) was performed to cluster the Keio strains into groups for GO enrichment analysis. A maximum distance cut-off (max distance: 30) was chosen based on visual inspection of the dendrogram, yielding 21 non-singleton clusters for GO enrichment analysis. Hierarchically clustered heatmaps were plotted using the *seaborn* module in Python.

#### Behavior monitoring and media supplementations

Sterile clear flat-bottom 6-well microplates (Corning Costar) were used for imaging. Bacteria were grown following the same protocol cited previously, and 30 µL of bacterial overnight liquid culture in LB was added at the center of the well containing NGM agar. Plates were then wrapped in wet tissues and incubated for 8 h at 25°C. For chemical supplementations, each chemical was dissolved in water or DMSO for the stock solution. Ascorbic acid (vitamin C, Sigma-Aldrich, A92902, 20 g/L), N-acetyl cysteine (NAC, Sigma-Aldrich, A7250, 20 g/L), paraquat (1,1′-dimethyl-4,4′-bipyridinium dichloride hydrate, Sigma-Aldrich, 36541, 0.5 and 1 mM), Fe_2_(SO_4_)_3_ (Sigma-Aldrich, 307718, 1 mM), shikimic acid (Sigma-Aldrich, S5375, 1 mM), gentisic acid sodium salt hydrate (Sigma-Aldrich, G5129, 1 mM), 2,3-dihydroxybenzoic acid (Sigma-Aldrich, 126209, 1 mM) were spotted on plates on top of the bacterial lawn 1 day prior to worm transfer.

### Bacterial growth assays and growth conditions

*E. coli* were cultured in LB, LB+kanamycin (50 mg/mL) for the deletion strains, LB+chloramphenicol (30 mg/L) for the overexpression strains, or LB+ampicillin (100 mg/L) for the RNAi strains. Strains were subsequently passaged on LB agar, then in LB broth with the corresponding antibiotics, and finally cultured once in antibiotic-free LB broth to minimize potential effects of residual antibiotics on *C. elegans* experiments. Liquid cultures were grown overnight (∼16 h) at 37°C with 200 rpm shaking. Bacterial strains of interest were transferred from stock plates into Nunc 96-well polystyrene round bottom plates containing 200 µL of LB supplemented with kanamycin (50 mg/mL) and grown overnight at 37°C. Using a 96-pin replicator, bacteria were transferred into a plate containing only 200 µL LB and incubated overnight at 37°C. Then 10 µL of each culture was transferred in Nunc 96-well round bottom plates containing 190 µL LB and placed in the BioTeK BioSpa 8 automated incubator (Agilent). The OD_600_ was recorded every 30 min for 24 h by the BioTek Citation 3 plate reader (Agilent).

### Bacterial mutant strain constructions

Double and triple gene deletions were generated using strains from the Keio Library. The kanamycin cassette was removed by using the plasmid pCP20 in *E. coli* BW25113 single mutant strains. This plasmid contains the yeast Flp recombinase gene, FLP, with chloramphenicol and ampicillin resistance genes, and its replication is temperature-sensitive ^56^. *E. coli* single mutant strains were transformed using the TSS enhanced chemical transformation protocol ^57^. Transformants were plated on chloramphenicol (30 mg/L) and incubated at 30°C overnight. Clones were selected and streaked on LB plates with no antibiotic selection and LB containing kanamycin (50 mg/L), and incubated at 30°C overnight. Kanamycin-sensitive clones were then streaked on LB with no selection, and incubated at 37°C overnight to stop the replication of the pCP20. Sensitive clones to kanamycin were confirmed by PCR for kanamycin cassette removal. Second and third mutations were introduced using another mutant from the Keio library and were lysed using P1 phage and transduced in the original kanamycin-sensitive *E. coli* strain. For loci separated by less than 100 kb, λ Red recombineering using the pKD46 plasmid was employed as an alternative strategy. Briefly, ∼60-bp primers were designed to include homology arms flanking the target gene and sequences complementary to the kanamycin resistance cassette. Using pKD4 as a template, PCR was performed to amplify the kanamycin resistance cassette with flanking homology arms. The resulting PCR products were electroporated into recipient strains expressing λ Red recombinase from pKD46 to replace the target gene.

We generated *E. coli* BW25113 compatible with RNAi experiments. For this, we used the phage P1 on *E. coli* K12 HT115 containing rnc14 :: DTn10 (RNAse III) and on *E. coli* CH1681 containing laczgA :: T7pol camFRT (RNA polymerase T7). Strains transduced with rnc and RNA polymerase T7 mutation were resistant to tetracyclin and cloramphenicol respectively. *E. coli* BW25113(DE3) and *E. coli* BW25113(DE3) Δ*fepD* were transformed with pL4440 plasmids encoding double-stranded RNA for RNAi of *C. elegans*. Mutations were confirmed by PCR using primers listed in the key resource table.

### Siderophore CAS assay

The CAS–MM9 and CAS–NGM agar used in this study were adapted and optimized based on previously described methods ^58^. Briefly, a 10× CAS reagent was prepared and mixed with either molten MM9 agar or NGM agar to generate the corresponding 1× CAS media. To prepare 100 mL of 10× CAS reagent, FeCl_3_ (Sigma-Aldrich) was dissolved in 10 mL of 10 mM HCl (Fisher Scientific) at a final concentration of either 1 mM or 100 mM to generate the iron(III) stock solution, corresponding to final Fe^3+^ concentrations of 10 µM or 1 mM in the completed CAS agar, respectively. The iron(III) solution was then mixed with 50 mL Chrome Azurol S solution (60.5 mg; Sigma-Aldrich). This iron(III)–CAS mixture was slowly added to 40 mL hexadecyltrimethylammonium bromide (HDTMA; 72.9 mg; Sigma-Aldrich) solution with gentle stirring to form the dark blue CAS complex. The resulting 10× CAS reagent was sterilized by filtration through a 0.22 µm membrane filter (Fisher Scientific). To prepare 400 mL of CAS–MM9 agar, 40 mL of 10× MM9 salts (per 100 mL: 0.3 g KH₂PO₄, 5 g NaCl, and 10 g NH₄Cl), 6 g Bacto agar, 6.048 g PIPES (Sigma-Aldrich), and 340 mL ddH₂O were combined, and the pH was adjusted to 6.8 using 10 M NaOH. Following autoclaving and cooling to ∼55°C, the following sterile supplements were added: 12 mL casamino acids, 8 mL 20% (w/v) glucose, 0.8 mL 1 M MgSO₄, and 40 µL 1 M CaCl_2_. The molten MM9 agar was then mixed with the 10× CAS reagent at a 9:1 ratio to generate 1× CAS–MM9 agar. To prepare CAS–NGM agar, molten NGM agar was similarly supplemented with 10× CAS reagent at a 9:1 ratio to generate 1× CAS–NGM agar. CAS–NGM and CAS–MM9 agar were poured into 9-cm Petri dishes (20 mL per plate). Overnight cultures of *E. coli* BW25113 and the indicated mutant strains were spotted onto CAS–NGM agar plates (10 µL per spot) or CAS–MM9 agar plates containing either 10 µM or 1 mM Fe³⁺, and incubated at 37°C for 48 h. Siderophore production was assessed by measuring the diameter of the halos formed by siderophore-mediated iron chelation using Fiji/ImageJ ^59^.

### Bacterial and nematode proteomics

#### Preparation of bacterial samples for proteomic analysis

Two proteomic experiments were performed: #1 with *E. coli* BW25113 wild-type, Δ*fepD::kan*, Δ*nuoC::kan*, Δ*atpB::kan*, and Δ*sdhD::kan*; and #2 with *E. coli* BW25113 wild-type, Δ*fepD::kan,* Δ*fepD*Δ*aceE::kan*, and Δ*aceE::kan*. Strains were seeded in LB broth overnight at 37°C. 9 cm NGM plates were seeded with 500 mL of overnight bacterial cultures and lawns were grown at 25°C for 8 h. 5 independent biological replicates were included per condition. Bacteria were collected from plates with PBS 1x buffer using a sterile glass scraper in diagenode tubes. Samples were centrifuged at 14000 rpm for 90 s at room temperature. The supernatant was removed and pellets were resuspended with 300 mL of lysis buffer (8 M urea, 20 mM HEPES pH 8). Samples were flash frozen in liquid nitrogen and kept on ice from this point onward. Pellets were then lysed via sonication for 5 min at 100% amplitude using a QSonica Q700 sonicator waterbath. Samples were centrifuged at 20000 g for 15 min at 4°C to separate the cellular debris and proteins. Supernatants containing the extracted protein were transferred to clean tubes, and protein concentrations were determined by the Quick Start Bradford protein assay (Bio-Rad) at 565 nm, with BSA used for standard curves.

#### Proteomic analysis with liquid chromatography-tandem mass spectrometry (LC-MS/MS) analysis

##### First proteomic analysis

Protein samples (50 µg/sample in 8 M urea) were processed using an in-solution digestion procedure. Briefly, samples were sequentially reduced and alkylated at room temperature and in the dark, to final concentrations of 5 mM dithiothreitol (DTT) and 15 mM 2-chloroacetamide (2-CAM), respectively. Samples were diluted 5-fold with 20 mM HEPES (pH 8.0), reducing the urea concentration to 1.5 M. This was followed by addition of 2 µg of trypsin (Serva, 37286). Samples were incubated overnight at 37°C. The digestion was stopped by acidification with 10% trifluoroacetic acid (TFA) to a final concentration of 0.5%, and protein digests were desalted using Glygen C18 spin tips (Glygen Corp, TT2C18.96). Tryptic peptides were eluted with 60% acetonitrile, 0.1% formic acid (FA), and dried by vacuum centrifugation. Dried tryptic digests were re-dissolved in 0.1% TFA and each sample injected at 1.5 µg on-column. LC-MS/MS analysis was performed using an Ultimate 3000 RSLC nano liquid chromatography system coupled to a Q-Exactive HF-X mass spectrometer via an EASY spray source (Thermo Scientific). For LC-MS/MS analysis re-dissolved protein digests were injected and loaded onto a trap column (Acclaim PepMap 100 C18, 100 μm × 2 cm) for desalting and concentrating at 8 μL/min in 2% acetonitrile, 0.1% TFA. Peptides were then eluted on-line to an analytical column (Acclaim Pepmap RSLC C18, 75 μm × 50 cm) at a flow rate of 250 nL/min. Peptides were separated using a 90 min gradient, 1-22% of buffer B for 60 min followed by 22-42% buffer B for another 30 min (composition of buffer B: 75% acetonitrile, 5% DMSO, 0.1% FA), with subsequent column conditioning and equilibration. Eluted peptides were analyzed by the mass spectrometer operating in positive polarity using a data-dependent acquisition mode. Ions for fragmentation were determined from an initial MS1 survey scan at 120000 resolution, followed by HCD (Higher Energy Collision Induced Dissociation) of the top 30 most abundant ions at 15000 resolution. MS1 and MS2 scan AGC targets were set to 3e6 and 5e4 for maximum injection times of 25 ms and 50 ms respectively. A survey scan m/z range of 375–1750 was used, normalised collision energy set to 27%, charge exclusion enabled with unassigned and +1 charge states rejected and a minimal AGC target of 8e3. Dynamic exclusion was set to 50 s.

##### Second proteomic analysis

Protein samples (100 µg/sample) were given to the proteomic platform. Samples were digested following a modified SP3 protocol ^60^, foregoing the second day peptide clean-up in favor of StageTip clean-up ^61^. Samples were analyzed by the CECAD Proteomics Facility on an Orbitrap Exploris 480 mass spectrometer coupled to a Vanquish Neo in reverse-flow flow setup (all Thermo Scientific). Samples were loaded onto a PepMap precolumn cartridge (#160434) with a flow of 15 µL/min eluent A (0.1% FA) before flushed onto an Aurora Ultimate 25 cm column (IonOpticks). Peptides were chromatographically separated with a flow rate of 300 nL/min on a linear gradient ranging from 4% B (0.1% FA in 80% acetonitrile) to 35% B in 77 min, followed by column wash with 98% B at 400 nL/min and re-equilibration to initial condition. MS1 scans were acquired from 399 m/z to 1001 m/z at 30 k resolution. Maximum injection time was set to 54 ms and the AGC target to 300%. MS2 scans ranged from 400 m/z to 1000 m/z and were acquired at 30k resolution with a maximum injection time of 54 ms, an AGC target of 100%, and were acquired in 30 x 20 m/z windows with an overlap of 1 m/z. All scans were stored as centroid. For the gas-phase fractionated library ^62^, a pool generated from all samples was analyzed in six individual runs covering the range from 400–1000 m/z in 100 m/z increments using identical LC settings as the samples. For each run, MS1 was acquired at 60k resolution with a maximum injection time of 98 ms and an AGC target of 100%. MS2 spectra were acquired at 30k resolution with a maximum injection time of 60 ms. Spectra were acquired in staggered 4 m/z windows, resulting in nominal 2 m/z windows after deconvolution using ProteoWizard ^63^.

#### Proteomic raw data processing

##### First proteomic analysis

Data was processed using the MaxQuant software platform (version 1.6.10.43), with database searches carried out with the Andromeda search engine against the UniProt *E. coli* database (version 20220310, number of entries: 4,402). A reverse decoy search approach was used at a 1% false discovery rate (FDR) for both peptide spectrum matches and protein groups. Search parameters included: maximum missed cleavages set to 3, fixed modification of cysteine carbamidomethylation and variable modifications of protein N-terminal acetylation, methionine oxidation, asparagine deamidation and N-terminal glutamine to pyro-glutamic acid. Label-free quantification was enabled with an LFQ minimum ratio count of 1. The ‘Match between runs’ function was used with match and alignment time limits of 0.7 and 20 min respectively.

##### Second proteomic analysis

The gas-phase fractionated library was built using DIA-NN (version 1.8.1) ^64^ and a SwissProt *E. coli* K12 database with settings matching acquisition parameters. DIA-NN was run with the additional command line prompts “—report-lib-info” and “—relaxed-prot-inf”. Further output settings were: filtered at 0.01 FDR; N-terminal methionine excision enabled; maximum number of missed cleavages =1; minimum peptide length =7; maximum peptide length =30; minimum precursor m/z =400; maximum precursor m/z =1000; and cysteine carbamidomethylation enabled as a fixed modification. Afterwards, DIA-NN output was further filtered on library q-value and global q-value ≤0.01 and at least two unique peptides per protein using R (version 4.1.3). Finally, LFQ values calculated using the DIA-NN R-package. Afterwards, analysis of results was performed in Perseus (version 1.6.15) ^65^ by filtering for data completeness in replicate groups, FDR-controlled *t*-tests, and 1D enrichment analysis.

#### Thermal Proteome Profiling (TPP) - Temperature gradient precipitation

Thermal proteome profiling was performed using a modified protocol based on ^66,67^. Briefly, proteins were extracted from 1-day-old adult N2 worms in HMP buffer (HEPES-KOH pH 7.5, 10 mM MgCl₂, cOmplete EDTA-free protease inhibitor (Roche), 0.4% NP-40) and quantified using a Bradford assay (Thermo Fisher Scientific). Enterobactin (Sigma-Aldrich) and 2,3-dihydroxybenzoic acid (DHB; Sigma-Aldrich) stock solutions were prepared in HMP buffer supplemented with 10% DMSO, while the vehicle control consisted of HMP buffer with 10% DMSO. Ligand enterobactin, DHB or control solution was added to the protein samples at a 1:20 ratio, resulting in final concentrations of ∼2 mg/mL protein, 0.1 mM ligand, and 0.5% DMSO in LoBind tubes. Samples were incubated at 25°C for 10 min with shaking at 600 rpm. Following incubation, 110 µL aliquots in PCR tubes (in triplicate) were subjected to a temperature gradient (25°C, 33°C, 37.9°C, 43.1°C, 47°C, 50.9°C, 55.1°C, 59°C, 63.9°C, and 69.1°C) using a thermocycler for 10 min, then cooled on ice for 2 min and centrifuged at 18,000 × g for 25 min at 4°C. A total of 90 µL of the supernatant was transferred to fresh PCR tubes, and protein concentrations for each temperature were determined using the Bradford assay (Thermo Fisher Scientific). Samples were then spiked with 0.8 µg BSA in 80 µL (1 µg/100 µL) as an internal standard, flash-frozen in liquid nitrogen, and lyophilized overnight. Pellets were resuspended in individual volumes of SDT buffer (4% SDS, 0.1 M DTT, 0.1 M Tris-HCl, pH 7.6) to a final concentration of 1 µg/µL, followed by protein digestion and sample preparation for proteome analysis via shotgun mass spectrometry as previously described in detail ^68^.

#### Thermal proteome profiling data analysis

The MS raw files were processed using MaxQuant software (version 2.4.13.0 ^69^), based on the *Caenorhabditis elegans* reference proteome obtained from UniProt (UP000001940; downloaded on 2026-03-11). For the generation of melting curve plots, statistical analysis, and identification of interacting candidates, the R script ‘Melting Curve Profiler (MeCuP)’ was used (https://github.com/andreaschrader/MeCuP) ^68^. Protein abundances (iBAQ values from MaxQuant) were normalized to the internal standard (BSA spike-in). Additionally, a starting point (SP) mean normalization was applied at the lowest temperature (25°C) separately for control and ligand-treated samples. Melting points (*T_m_*_50_) were determined from LOESS-smoothed curves as the temperature at which the soluble fraction reaches 50%. Protein melting curves that show non-overlapping profiles between control and treatment conditions indicate a potential interaction between the protein and the ligand. Differential protein analysis was performed in R (RStudio environment), and functional enrichment and protein interaction network analyses were conducted using the STRINGdb package^70^. The mass spectrometry proteomics data generated in this study have been deposited in the PRIDE partner repository via the ProteomeXchange Consortium with the dataset identifiers, PXD075235, PXD075268 and PXD078857, respectively.

### Computational modeling of human glutaryl-CoA dehydrogenase binding by enterobactin

The interaction of enterobactin with human glutaryl-CoA dehydrogenase was investigated using the web-based molecular docking platform SwissDock employing the Attracting Cavities 2.0 docking protocol^22^. The crystal structure of human glutaryl-CoA dehydrogenase (PDB ID: 1SIQ) was retrieved from the Protein Data Bank and used as the receptor structure without additional target preparation. The docking region was centered close to the FAD cofactor to probe potential ligand interactions in the catalytically relevant region of the enzyme. The ligand structure was generated from the corresponding SMILES representation and submitted to SwissDock without additional geometry optimization. The following docking parameters were eventually chosen as optimal, with regard to biochemically reasonable poses and favorable AC docking score values:

Ligand: OC1=CC=CC(C(=O)NC2COC(=O)C(COC(=O)C(COC2=O)NC(=O)C2=C(O)C(O)=CC=C2)NC(=O)C2=C(O)C(O)=CC=C2)=C1O; Target: human glutaryl-CoA dehydrogenase + FAD (PDB ID: 1SIQ); Sampling exhaustivity =90; Cavity prioritization =70; Random Initial Conditions =1; Box size = 25 Å, 25 Å, 25 Å; Box center = –18.0 Å, 45.0 Å, –47.0 Å.

The docking pose with the most favorable AC docking score was selected for further analysis. The docking results associated with the aforementioned parameters are publicly available on SwissDock (https://www.swissdock.ch/results.php?job=65218376).

### Human cell western blot analysis

HCT116 cells were seeded in 12-well plates and cultured overnight to reach ∼80–90% confluency. Cells were then treated with 100 μM enterobactin, 100 μM actinonin (Sigma-Aldrich), or vehicle control (1% DMSO) in 400 μL RPMI 1640 medium supplemented with 1% FBS and 1% penicillin/streptomycin for 12 h or 24 h. Three independent biological replicates were included for each condition. Following treatment, cells were washed once with ice-cold Dulbecco’s phosphate-buffered saline (DPBS, Thermo Fisher Scientific) and lyzed in RIPA buffer supplemented with cOmplete Protease Inhibitor Cocktail (Sigma-Aldrich) and PhosSTOP phosphatase inhibitor (Roche). Lysates were incubated at 4°C for 30 min with gentle rotation and subsequently sonicated using a Qsonica Q800R3 sonicator (40% amplitude, 10 cycles of 3 s on and 3 s off). Samples were clarified by centrifugation at 14,000 × g for 35 min at 4°C, and protein concentrations were determined using a Pierce BCA Protein Assay Kit (Thermo Fisher Scientific). Equal amounts of protein were mixed with 6× Laemmli sample buffer (Thermo Fisher Scientific), heated at 95°C for 3 min, and separated on 4–20% Criterion TGX Precast Midi Protein Gels (Bio-Rad). Precision Plus Protein Dual Color Standards (Bio-Rad) were used as molecular weight markers. Electrophoresis was performed at 90 V for 30 min followed by 130 V for 1 h. Proteins were transferred onto polyvinylidene difluoride (PVDF) membranes using Trans-Blot Turbo PVDF Transfer Packs (Bio-Rad). Membranes were blocked in either 5% skim milk (Millipore) or 3% fish gelatin (Sigma-Aldrich), depending on the primary antibody used, and incubated overnight at 4°C with primary antibodies diluted 1:1,000 in the corresponding blocking buffer containing 0.02% sodium azide in PBST or TBST. The following primary antibodies were used: α-tubulin (Cell Signaling Technology, Cat# 2144), eIF2α (Abcam, Cat# ab26197), phospho-eIF2α (Abcam, Cat# ab32157), and ALDH18A1 (Proteintech, Cat# 17791-1-AP). Following washing, membranes were incubated with horseradish peroxidase (HRP)-conjugated goat anti-mouse IgG (Fc-specific) secondary antibody (Sigma-Aldrich, Cat# A0168) or HRP-conjugated anti-rabbit IgG secondary antibody (Cell Signaling Technology, Cat# 7074) at a dilution of 1:5,000, as appropriate. Immunoreactive bands were visualized using Clarity Western ECL Substrate (Bio-Rad, Cat# 1705061) according to the manufacturer’s instructions.

### Bacterial metabolomics

Levels of intermediates (organic acids, sugar phosphates) from glycolysis and the TCA cycle in *E. coli* were determined by Anion-Exchange Chromatography coupled to Electrospray Ionization High-Resolution Mass Spectrometry (IC-ESI-HRMS) using previously described procedures ^71,72^ with some modifications: BW, Δ*fepD::kan*, Δ*sdhD::kan* were cultured in NGM liquid for 16 h at 25°C, 200 rpm. Liquid cultures were split in two, one for the metabolomic analysis and one for the protein content. Pellets for metabolic analysis were suspended in 350 µL of ice-cold methanol/acetonitrile/water 5:3:2 (v/v/v) using the Precellys 24 homogenizer (Peqlab) at 6,400 rpm twice for 10 s with a 5 min pause. 150 µL of homogenate were diluted with 150 µL of methanol/acetonitrile/water 5:3:2 (v/v/v) and 25 µL of a mixture of isotope-labeled internal standards in Milli-Q water (5 µM ^13^C_6_-D-glucose-6-phosphate (^13^C_6_-G6P) and 5 µM *d*_4_-succinic acid (*d*_4_-SUC, Eurisotop) were added. The samples were extracted using a pre-cooled ThermoMixer (Eppendorf) at 4°C and 900 rpm for 20 min. After centrifugation (16,100 rcf, 20 min, 4°C), the supernatants were dried under a stream of nitrogen, and the residues were resolved in 100 µL of Milli-Q water. After mixing and centrifugation, 80 µL of supernatant were transferred to autoinjector vials and measured immediately.

IC-HRMS analysis was performed using a Dionex Integrion RFIC system (Thermo Scientific) equipped with a Dionex IonPac AS11-HC column (2 mm × 250 mm, 4 μm particle size, Thermo Scientific) and a Dionex IonPac AG11-HC guard column (2 mm × 50 mm, 4 μm, Thermo Scientific) and coupled to a Q Exactive HF quadrupole-orbitrap mass spectrometer (Thermo Scientific). 5 µL of sample were injected using a Dionex AS-AP at 10°C. The IC was operated at a flow rate of 0.38 mL/min with a potassium hydroxide gradient which was produced by an eluent generator with a potassium hydroxide cartridge and Milli-Q water. The gradient started with 10 mM KOH over 3 min, 10−50 mM from 3 to 12 min, 50-100 mM from 12 to 19 min, held at 100 mM from 19 to 25 min, and re-equilibrated at 10 mM for 3 min. The total run time was 28 min. An Dionex ADRS 600, 2 mm suppressor was operated with 95 mA, and methanol was used to produce a make-up flow at a flow rate of 0.15 mL/min.

The mass spectrometer was operated in negative ion mode. Full MS scans in the range of m/z 60-900 were acquired with a resolution of 120,000, an Automatic Gain Control (AGC) target value of 1 x 10^6^ and a maximum injection time (IT) of 240 ms. Spectrum data were collected in the centroid mode. The ESI source was operated with flow rates for sheath gas, auxiliary gas, and sweep gas of 50, 14 and 3, respectively. The spray voltage setting was 2.75 kV, the capillary temperature 230°C, the S-lens RF level 45, and the auxiliary gas heater temperature 380°C ^71,72^.

The exact m/z traces of the internal standards and endogenous organic acids and sugar phosphates were extracted and integrated using TraceFinder software (version 5.1; Thermo Scientific). Endogenous metabolites were quantified by normalizing their peak areas to those of the internal standards: ^13^C_6_-G6P was used as internal standard for sugar phosphates and *d*_4_-SUC for organic acids. These peak area ratios were normalized to the protein content of the homogenate determined in an extra sample for each group, which was incubated under the same conditions.

### Nematode bulk RNA-seq

1-day-old adult N2 worms fed on BW25113 were synchronized to L1 stage by standard hypochlorite bleaching and transferred onto 3-day bacteria-seeded NGM plates, at a density of 300 worms per plate. For the Δ*fepD*::kan and Δ*fepD*Δ*entF*::kan conditions, 10 µM 2,3-dihydroxybenzoic acid (DHB) was added to the bacterial lawns 1 day prior to L1 larval transfer to enhance enterobactin biosynthesis. Worms were collected 24 h after reaching the L4 stage, corresponding to day 1 adulthood, with minor adjustments in collection time depending on developmental rate under different conditions.

Approximately 1,800 worms were collected per biological replicate by washing three times with 10 mL PBS, followed by collection in 10 mL RNase-free water on ice. Worm pellets were transferred to 2 mL RNase-free tubes and washed three additional times with 1 mL RNase free water (Thermo Fisher Scientific) on ice. After removal of residual liquid, 350 µL TRIzol reagent was added together with ceramic beads (Avantor, Inc., USA). Samples were flash-frozen in liquid nitrogen and homogenized at 6500 rpm for 30 s twice using a bead homogenizer (Avantor, Inc., USA). Lysates were clarified by centrifugation at 14000 rpm for 20 min at 4°C, and total RNA was extracted using the Direct-zol RNA Miniprep kit (Zymo Research). RNA concentration was measured using a NanoDrop 2000 spectrophotometer (Thermo Fisher Scientific) and Qubit RNA assay kit (Thermo Fisher Scientific). RNA integrity was assessed using an Agilent 2100 Bioanalyzer (Agilent Technologies). RNA-seq libraries were made from 200ng Total RNA using the Watchmaker Genomics Ltd mRNA Library Prep and RNA Library Prep Kits (New England Biolabs) according to manufacturer’s instructions. Libraries were dual indexed using IDT xGen Stubby Adapter-UDI Primers. Library quality and quantity was checked using the Agilent 4200 Tapestation and Qubit fluorometer respectively and then pooled in an equimolar manner and run on an Illumina NextSeq2000 sequencer, generating a minimum of 20 million paired-end 60 bp reads per sample.

Transcript quantification was performed using tximport ^73^, differential gene expression analysis was conducted with DESeq2 ^74^, and downstream analyses and visualization were carried out using tidyverse, ComplexHeatmap ^75^, and WormCat ^76^. Gene annotations were retrieved from Ensembl using ensembldb ^77^ and Bioconductor AnnotationHub ^78^. RNA-seq data have been deposited in GEO under accession number <u>GSE329272</u>.

### Nematode Fluorescence Measurements

#### UPR^mt^ measurements

To measure mitochondrial UPR (UPR^mt^) in *C. elegans*, we used a crossed strain of SJ4100 and WMB1143 with the genotype P*hsp-6*::GFP;*eft-3p*::Mcherry. NGM plates were seeded with bacteria 3 days before L1 seeding. A minimum of 200 L1 worms were transferred per replicate per condition, and repeated three times independently, around 20 worms from each plate were anesthetized with 2% levamisole on NGM plates. Worms were imaged under a 40x objective using a Zeiss Axio Zoom V16 microscope system equipped with an AxioCam MRm camera operated by Zen 2 software (Zeiss). Either the GFP filterset (excitation: 450-490 nm; emission: 500-550 nm) or the mCherry filterset (excitation: 559-585 nm; emission: 600-690 nm) was used. All images were exported in CZI format and fluorescence levels were quantified using ImageJ. The fluorescence intensity of worms was calculated as the pixel density of the entire cross-sectional area occupied by worms from which the pixel density of the background had been subtracted. 3 independent trials were carried out with a minimum of 8 worms imaged per condition per trial. The fluorescence intensity was calculated automatically by setting a minimum threshold intensity that excluded the background. 3-6 independent biological replicates were included per condition. Statistical analysis was done by using a one-way ANOVA with multiple comparisons (Tukey’s multiple comparison test) with the software GraphPad Prism (version 8.4.0).

#### Cathecol-methyl transferase measurements

To assess catechol-methyltransferase expression levels in *C. elegans*, the reporter strain CER554 (*comt-4*p::GFP::H2B) was used, in which the endogenous *comt-4* locus, encoding a catechol-O-methyltransferase, is replaced by a GFP::H2B reporter under the control of the native *comt-4* promoter. Synchronized L1 larvae were transferred onto NGM plates seeded with wild-type *E. coli* BW25113 or the indicated mutant strains and cultured until the day-1 adult stage. A minimum of 200 L1 worms were transferred per biological replicate for each condition, and the experiment was independently repeated three times. Approximately 20 worms were collected from each plate and anesthetized with 2% levamisole on NGM plates for imaging. Worms were imaged under a 40× objective using a Zeiss Axio Zoom V16 microscope system equipped with an AxioCam MRm camera and operated using Zen 2 software (Zeiss), with a GFP filter set (excitation: 450-490 nm; emission: 500-550 nm). As both the proportion of worms exhibiting inducible *comt-4* reporter expression and the reporter signal intensity varied between conditions, two quantitative parameters were assessed: (1) the percentage of worms showing detectable *comt-4* reporter activation, and (2) the maximum fluorescence intensity of GFP-positive reporter puncta in individual worms following background subtraction. Images were exported in CZI format and analyzed using Fiji/ImageJ.

#### Blue fluorescence measurements

To measure the intestinal autofluorescence of *C. elegans*, the synchronized L1 worms were transferred onto NGM plates seeded with bacteria. Autofluorescence was assessed on the first day of adulthood. A minimum of 200 L1 worms were transferred per replicate per condition, and repeated three times independently, around 20 worms from each plate were anesthetized with 2% levamisole on NGM plates. Worms were imaged under a 40× objective using a Zeiss Axio Zoom V16 microscope system equipped with an AxioCam MRm camera and operated using Zen 2 software (Zeiss), with a DAPI filter set (excitation: 335-383 nm; emission: 420-470 nm). Images were exported in CZI format, and fluorescence intensity was quantified using Fiji/ImageJ. Whole-worm fluorescence intensity was calculated as the sum of pixel intensities across the entire worm area after subtraction of background fluorescence. Statistical analysis was performed using one-way ANOVA followed by Tukey’s multiple comparison test in GraphPad Prism (version 8.4.0).

#### ROS fluorescence measurements

To assess intestinal ROS-associated fluorescence of *C. elegans*, we used an intestinal redox/oxidative stress reporter line HCX1537 ^79^. Synchronized L1 worms were transferred onto NGM plates seeded with wild-type *E. coli* BW25113 or Δ*fepD* mutant bacteria, and cultured until the day-1 adult stage. A minimum of 200 L1 worms were transferred per replicate per condition, and repeated four times independently. Approximately 20 worms from each plate were anesthetized with 2% levamisole on NGM plates. Worms were imaged under a 40× objective using a Zeiss Axio Zoom V16 microscope system equipped with an AxioCam MRm camera and operated using Zen 2 software (Zeiss), with an mCherry filterset (excitation: 559-585 nm; emission: 600-690 nm). Images were exported in CZI format and analyzed using Fiji/ImageJ. 4 independent trials were carried out with a minimum of 8 worms imaged per condition. Whole-worm fluorescence intensity was calculated as the sum of pixel intensities across the entire worm area after subtraction of background fluorescence. Statistical analysis was performed using a student *t*-test in GraphPad Prism (version 8.4.0).

### Nematode lifespan assay

Bacteria were grown overnight in LB broth at 37°C, 200 rpm and seeded onto NGM plates (100 µL per plate) 3 days prior to worm transfer. N2 and mutant worms were synchronized by standard hypochlorite bleaching and arrested at the L1 stage, then transferred onto 3-day-old bacterial lawns. Worms were transferred again at the L4 stage to fresh plates under the same conditions to define day 0 of adulthood. Worms were subsequently transferred daily for the following 5 days and again on day 7 to minimize confounding effects from progeny. Dead and censored worms were scored every 2 days. Approximately 120 worms were analyzed per condition starting from day 0 of adulthood. For lifespan assays on RNAi bacteria (BW25113 (DE3) and Δ*fepD* (DE3)), bacterial overnight cultures were seeded onto NGM agar supplemented with ampicillin (100 µg/mL) and IPTG (final concentration 1 mM) at 200 µL per plate. N2 worms synchronized by bleaching were transferred onto 1-day-old RNAi bacterial lawns. Worms were transferred at the L4 stage to fresh RNAi plates to define day 0 of adulthood and were subsequently transferred daily until day 7 to minimize the effects of progeny. Dead and censored worms were scored every 2 days. Approximately 120 worms were analyzed per condition starting from day 0 of adulthood. Survival curves were generated using Kaplan–Meier analysis and plotted using GraphPad Prism (version 11).

### Nematode body bends assay

To quantify the number of body bends, a nematode was transferred onto a 35 mm NGM plate without bacteria, and allowed to crawl for 1 min to remove any excess bacteria, after which the body bends were scored for 1 min. One body bend corresponds to the movement of the head region to the opposite direction. The assay was performed on day 1, day 5 and day 10 adults, using 10 animals per replicate per condition, and repeated three times independently.

### Nematode muscle fiber morphology and quantification

Day 1 and 15 adults of *raIs5* [*myo-3p::GFP::myo-3 + rol-6(su1006)*] were mounted on 2% agarose pads in a drop of 50 mM sodium azide, and imaged using the TCS SP8 gSTED 3X microscope (Leica Microsystems). To visualize all the muscle fibers in a single plane, we used Leica Application Suite X (LAS X) software to convert the Z-stacks into 2D images by applying maximum intensity projection, and these were then cropped in Image J (Version 1.54t) to be used for a semi-quantitative analysis of muscle morphology. To that end, we defined four distinctive categories: 1. normal = smooth, parallel, and continuous muscle fibers without gaps or breaks; 2. mild = slight waviness and minor disorganization in fiber alignment; 3. moderate = clear disruptions in fiber alignment and structure, including multiple gaps and clumping; and 4. severe = major loss of organized muscle structure, with severely fragmented, misaligned fibers ^80^. The blinded visual scoring was done on the level of the individual muscle cells, using the software Blinder (Solibyte Solutions, Durham, NC), as previously described ^81^. For each experimental condition, ∼10 animals were used per replicate, and the experiment was repeated three times independently.

### Nematode development assay

N2 wild-type and pdfr-1 knockout worms were synchronized to the L1 stage and cultured on NGM plates pre-seeded with 3-day-old bacterial lawns (BW25113 or mutant bacteria), with ∼100 worms per plate. After 48 h, developmental stages of individual worms on each plate were scored according to the following criteria: 0 = complete developmental arrest (arrested at the L1 stage); 1 = L2/L3-sized; 2= L4-sized; 3 = young adult-sized; 4 = fully developed, fertile adults with progeny.

