## Supplemental Figures 1 to 10 for "A Neuronally Active Bacterial Siderophore Manipulates Host Behavior and Longevity"

### EXTENDED DATA FIGURES

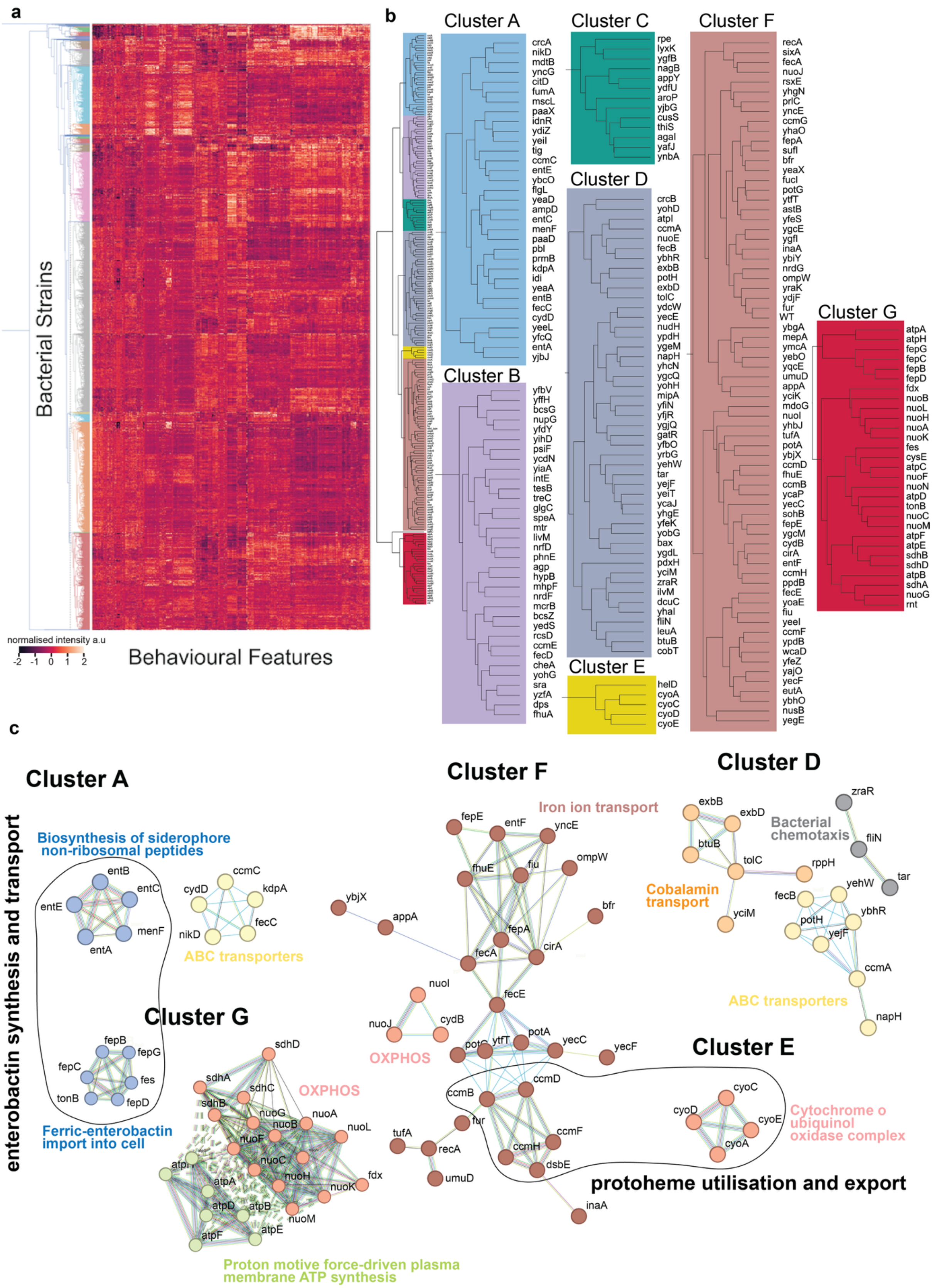


**Extended Data Figure 1. a,** Complete heatmap representation of the MH-BHTS dataset for *C. elegans* fed on each bacterial deletion mutant of the Keio library. **b,** Dendogram associated gene clusters based on behavioral features. **c,** STRINGdb k-means clustering with primary and/or secondary enrichment function for cluster number ≤3.

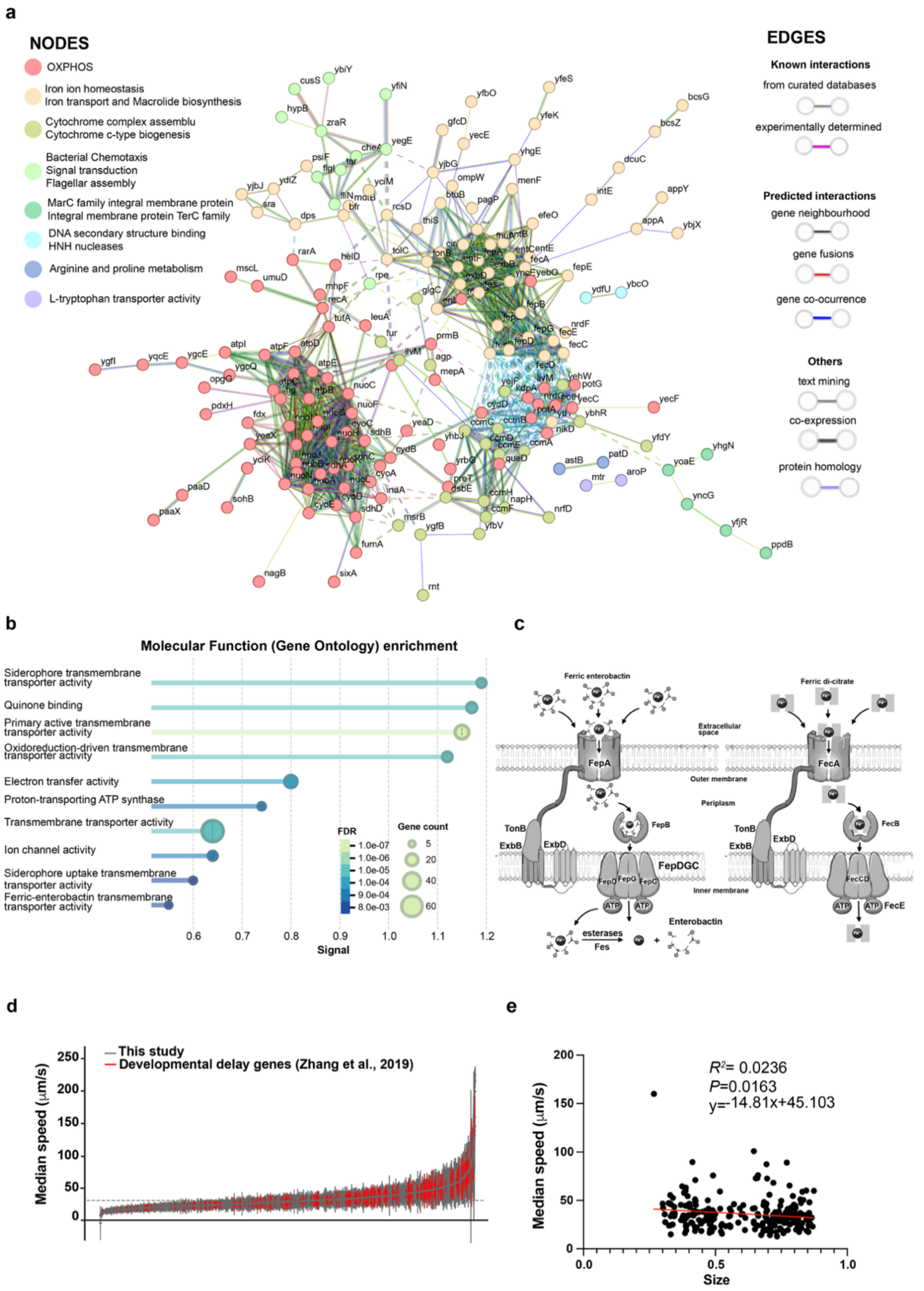


**Extended Figure Data 2. a,** STRINGdb k-means clustering with primary and/or secondary enrichment function for cluster number =8. **b,** Molecular function enrichment of significant clusters of genes altering *C. elegans* behavior sorted by strength (signal), False Discovery Rate (FDR), and gene count. **c,** Diagram of iron transport in *E. coli*. **d,** Comparison with Zhang *et al.* 2019 [21] of bacterial gene involvement in alteration of *C. elegans* median speed. **e,** Correlation analysis between worm size and locomotor speed (50th percentile).

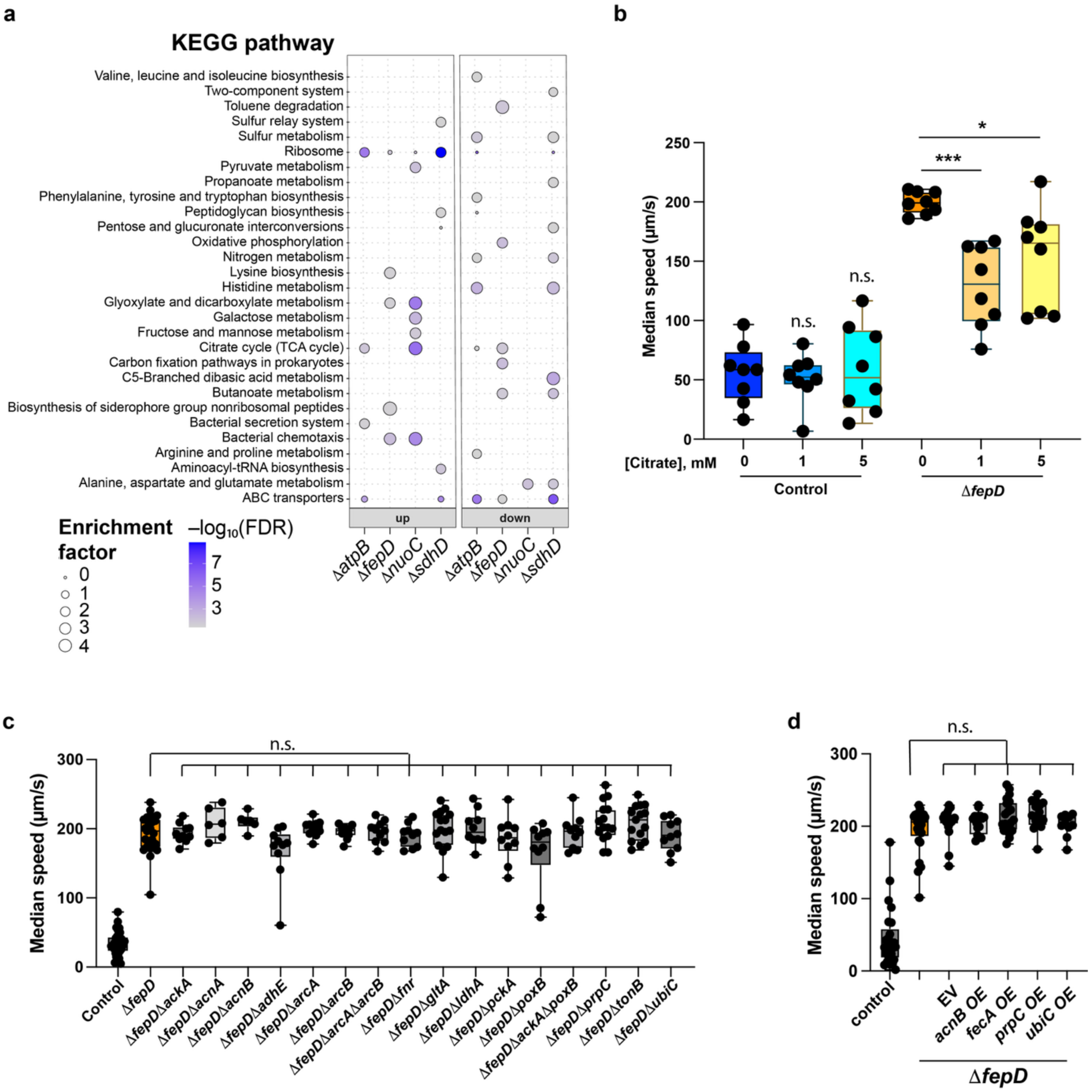


**Extended Data Figure 3. a,** KEGG pathways enriched of Δ*atpB*, Δ*fepD*, Δ*nuoC* and Δ*sdhD* from proteomics compared with BW25113 wild-type, data expressed in $-\log_{10}$ (FDR) and enrichment factor. **b,** Median speed of worms fed on control BW25113 and Δ*fepD* bacteria, with citrate supplementation (0, 1, 5 mM). Each dot represents the average of worms in one well (n=8 wells with 10 worms/well). **c**, Median speed of worms fed on Δ*fedD* bacteria with additional selected mutants affecting bacterial carbon metabolism and iron entrance regulation. Each dot represents the average of worms in one well (n=9-30 wells with 10 worms/well). **d,** Median speed of worms fed on control BW25113 and Δf*epD* bacteria with overexpression of *acnB, fecA, prpC* and *ubiC*. Each dot represents the average of worms in one well (n=10-25 wells with 10 worms/well). Stats correspond to comparison against the control (one-way ANOVA) represented as *p < 0.05, **p < 0.01, ***p < 0.001, n.s p > 0.05.

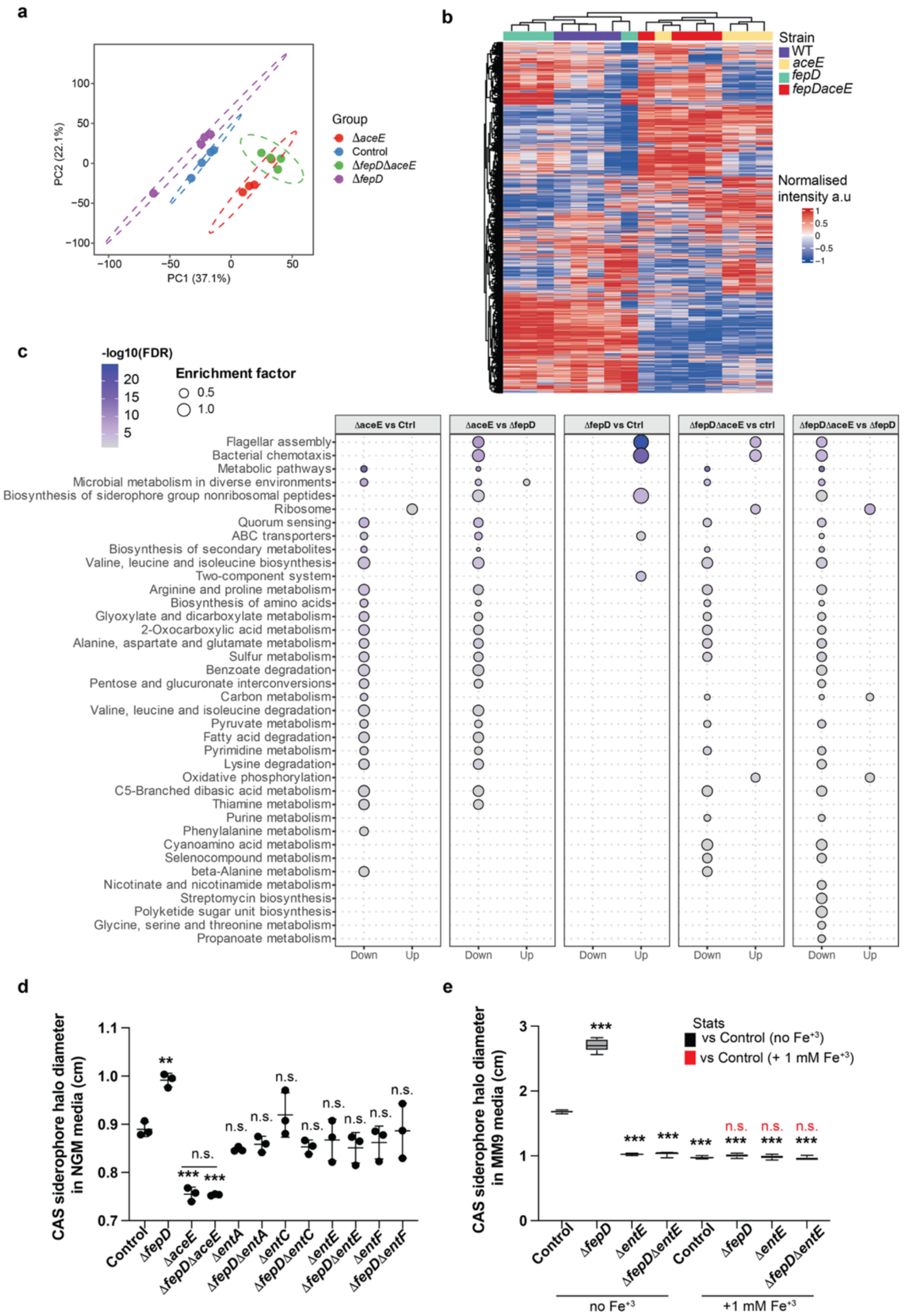


**Extended Data Figure 4. a,** Principal component analysis of proteomic profiles from control BW25113, Δ*aceE*, Δ*fepD* and Δ*fepD*Δ*aceE* bacteria. Each point represents an independent biological replicate with n=4. Dashed ellipses indicate group clustering. **b,** Unsupervised hierarchical clustering heatmap of proteomic profiles across BW25113, Δ*aceE*, Δ*fepD* and Δ*fepD*Δ*aceE* bacteria. Rows represent proteins and columns represent biological replicates with n=4. Protein intensities were row-wise z-score normalized and trimmed to the indicated range for visualization. Red indicates relatively higher abundance and blue indicates relatively lower abundance. **c,** Comparison of KEGG pathway enrichments between control BW25113, Δ*aceE*, Δ*fepD* and Δ*fepD*Δ*aceE* bacteria. Dot size represents the enrichment factor, and color intensity indicates statistical significance ($-\log_{10}$ FDR). **d,** Siderophore halo diameter in NGM for control BW25113, Δ*aceE*, Δ*fepD,* Δ*entA,* Δ*entC,* Δ*entE,* Δ*entF* single and double bacterial mutants, Δ*fepD*Δ*aceE* and Δ*fepD*Δ*entA/C/E/F.* Each dot represents one biological replicate with n=3. **e,** Siderophore halo diameter in NGM for control BW25113, Δ*fepD,* Δ*entE* and double mutant Δ*fepD*Δ*entE* bacteria in the presence or absence of 1 mM of $Fe^{3+}$. Each dot represents one biological replicate with n=3. Stats correspond to comparison against the control (one-way ANOVA) and between nested conditions (two-way ANOVA), represented as *$p < 0.05$, **$p < 0.01$, ***$p < 0.001$, n.s. $p > 0.05$.

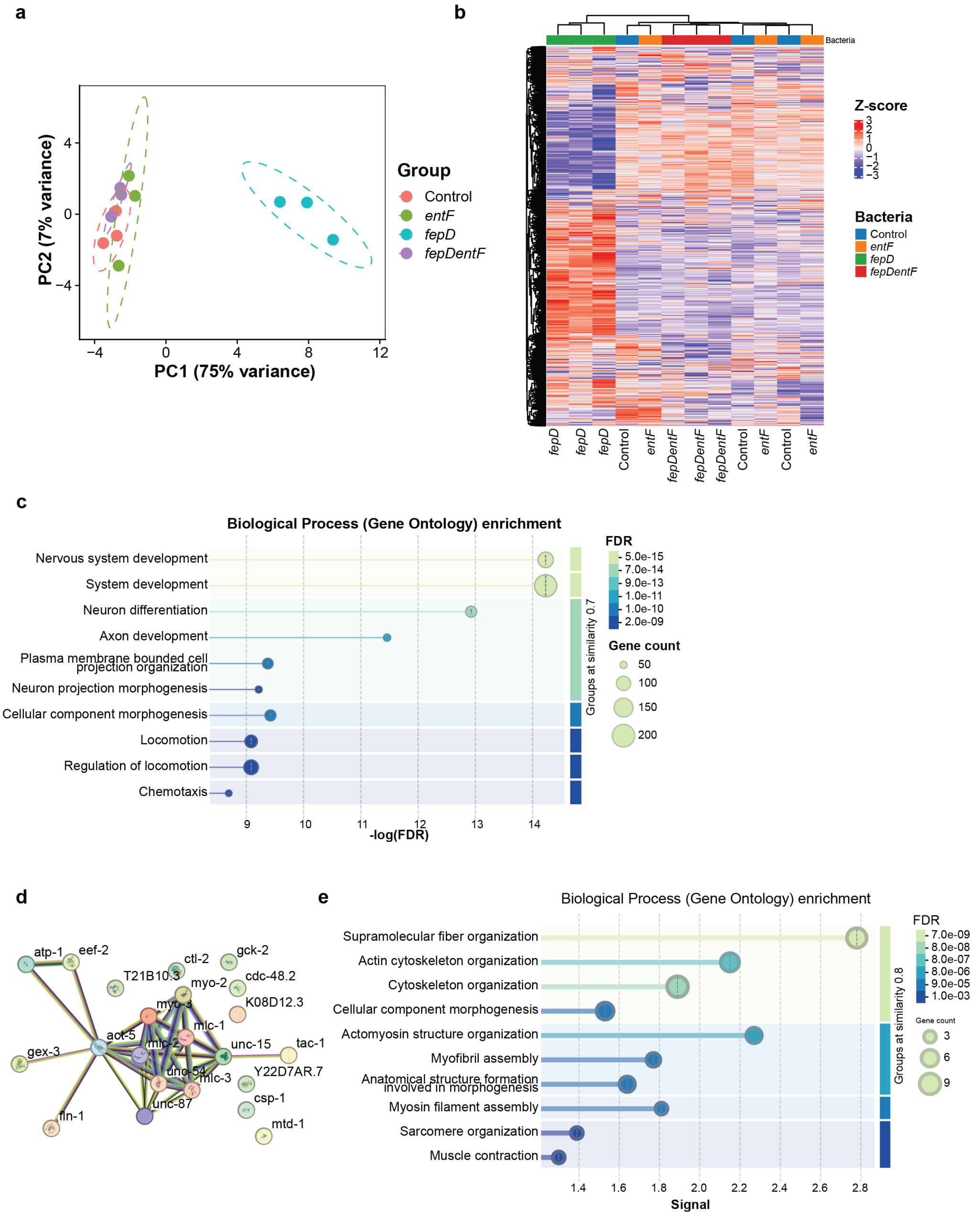


**Extended Data Figure 5. a,** Principal component analysis of RNA-seq profiles from N2 worms fed on control BW25113, Δ*entF*, Δ*fepD* or Δ*fepD*Δ*entF* bacteria. Each point represents an independent biological replicate with n=3. Dashed ellipses indicate group clustering. **b,** Unsupervised hierarchical clustering heatmap of the top 5,000 differentially expressed genes from RNA-seq profiles of the BW25113, Δ*fepD*, Δ*entF*, and Δ*fepD*Δ*entF* conditions. Rows represent individual genes, and columns represent biological replicates (n=3 per condition). Gene expression values were row-wise z-score normalized and trimmed to the indicated range for visualization. Red indicates relatively higher expression, whereas blue indicates relatively lower expression. **c,** FDR-ranked biological process enrichment for Δ*fepD* compared to the control BW25113. Dot size represents the gene count, and color intensity indicates statistical significance (FDR). **d,** Results of the RNA-seq analysis comparing N2 fed on control BW25113 and Δ*fepD* bacteria. STRINGdb network highlights a strong enrichment of muscle- and cytoskeleton-associated genes. **e**, FDR-ranked biological process enrichment for Δ*fepD* compared to the control BW25113. Dot size represents the gene count, and color intensity indicates statistical significance (FDR).

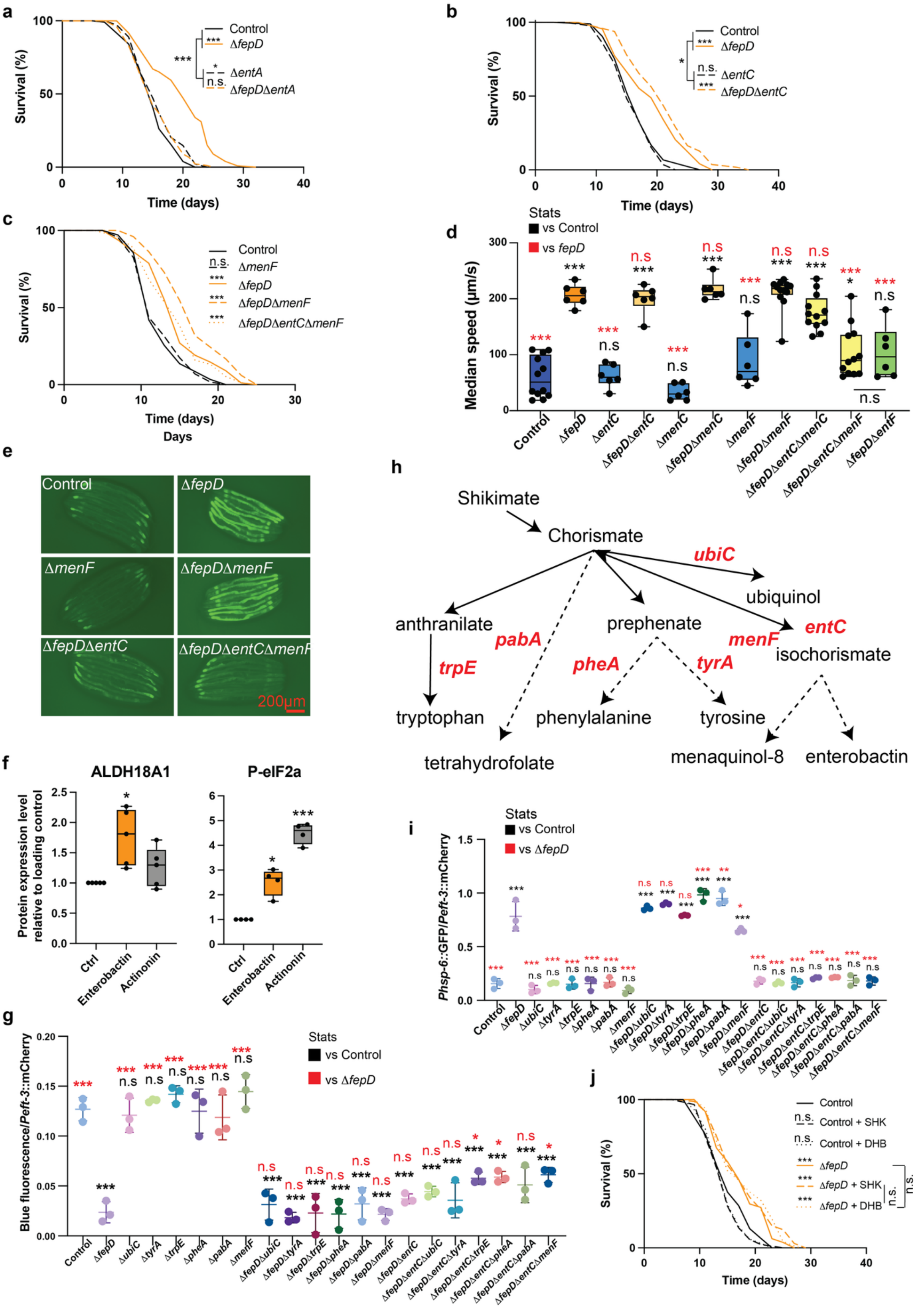


**Extended Data Figure 6. a-c,** Lifespans of N2 worms fed on control BW25113 or Δ*fepD* bacteria, with the following mutants: Δ*entA*, Δ*fepD*Δ*entA* (n=116-122, 2 independently run replicates) (**a**); Δ*entC*, Δ*fepD*Δ*entC* (n=110-129, 2 independently run replicates) (**b**); Δ*menF,* Δ*fepD*Δ*mentF,*

Δ*fepD*Δ*entC*Δ*mentF* (n=87-120) (**c**). **d,** Median speed of worms fed on control BW25113 or the indicated bacterial mutants. Each dot represents the average of worms in one well (n=6-12 wells with 10 worms/well). Statistical significance was determined relative to the control (black) or Δ*fepD* (red) conditions. **e,** Microscopy images of the reporter for *hsp-6* expression of worms fed on control BW25113, Δ*fepD,* Δ*menF,* Δ*fepD*Δ*menF,* Δ*fepD*Δ*entC* or Δ*fepD*Δ*entC*Δ*menF* bacteria. **f,** Protein level quantification of ALDH18A1 and elF2a in HCT116 cells treated with 100 μM enterobactin or actinonin, relative to tubulin as a loading control. **g,** Normalized age-associated intestinal blue autofluorescence over mCherry in N2 worms fed on control BW25113 or the indicated bacterial mutants. Each dot represents an average of 15 worms with n=3. **h,** Shikimate catabolic reactions in *E. coli*. Enzyme genes are shown in red. **i,** Normalized brightness of the worm reporter for *hsp-6* expression as a ratio of GFP over mCherry measured in N2 fed control BW25113 or the indicated bacterial mutants. Statistical significance was determined relative to control (black) or Δ*fepD* (red) conditions. Each dot represents an average of 15 worms with n=3. **j,** Lifespans of N2 worms fed on control BW25113 or Δ*fepD* bacteria, treated with 10 μM shikimic acid (SHK) or 10 μM 2,3-dihydroxybenzoic acid (DHB) (n=90-117). Lifespans were monitored with n=100. Microscopy images were analyzed with ImageJ. Statistics were performed against the control or Δ*fepD* mutant condition (one-way ANOVA, or Log-Rank for survival curves), between nested conditions (Cox Proportional Hazards for survival curves), represented as $^*p < 0.05$, $^{**}p < 0.01$, $^{***}p < 0.001$, n.s. $p > 0.05$.

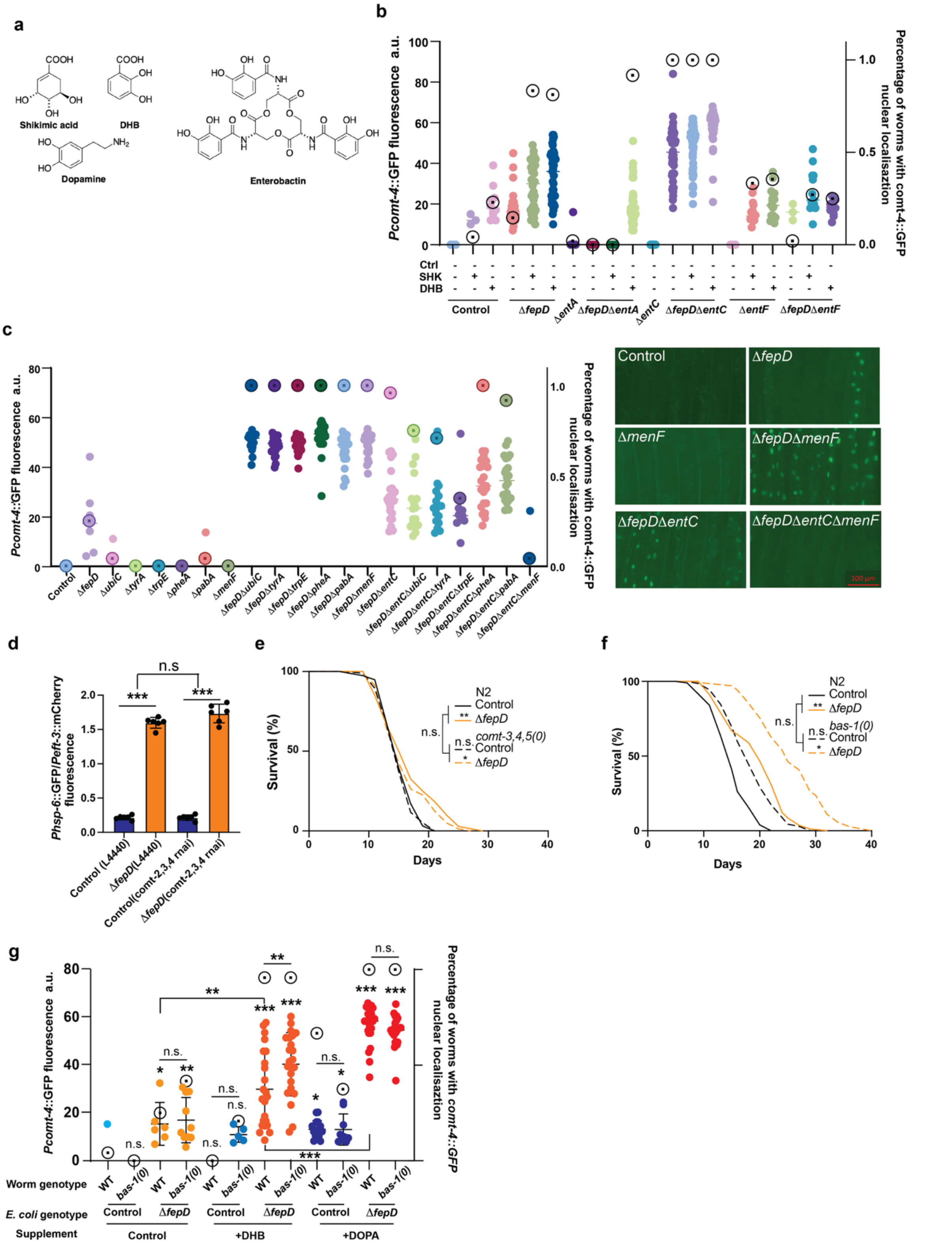

**Extended Data Figure 7. a,** Chemical structures of the catechol derivatives shikimate and 2,3-dihydroxybenzoic acid (DHB), key intermediates in enterobactin biosynthesis, alongside enterobactin itself and the neurotransmitter dopamine. **b,** *Pcomt-4*::GFP reporter response to enterobactin biosynthesis pathway perturbation. Worms were exposed to *E. coli* enterobactin synthesis/pathway-related knockout mutants, supplemented with or without shikimate (SHK, 10 µM) or 2,3-dihydroxybenzoic acid (DHB, 10 µM). The left y-axis shows the maximum *Pcomt-4*::GFP fluorescence intensity (a.u.), with each dot representing one worm. The right y-axis shows the fraction of worms exhibiting nuclear *Pcomt-4*::GFP localization. Open circles with central dots indicate values corresponding to the right y-axis. **c,** *Pcomt-4*::GFP reporter response to *E. coli* mutants affecting chorismate-derived enterobactin biosynthesis, with representative fluorescence images showing nuclear *Pcomt-4*::GFP localization under selected conditions. **d,** Normalized *hsp-6* reporter intensity in N2 worms, quantified as the GFP/mCherry fluorescence ratio, after feeding RNAi-mediated knockdown of catechol-O-methyltransferase-related genes *(comt-2, comt-3,* and *comt-4*). **e,** Lifespan analysis of N2 and *comt-3*, *comt-4*, and *comt-5* knockout worms fed on control BW25113 or Δ*fepD E. coli* (n=91-120, 2 independently run replicates). **f,** Lifespan analysis of N2 and the dopamine/serotonin biosynthetic gene *bas-1* knockout worms fed on control BW25113 or Δ*fepD E. coli* (n=94-124, 2 independently run replicates). **g,** *Pcomt-4*::GFP reporter activity in wild-type N2 and *bas-1* mutant worms fed control BW25113 or Δ*fepD E. coli*, supplemented with or without DHB (10 μM) or DOPA (dopamine hydrochloride, 10 mM). Statistics were performed against the control or Δ*fepD* mutant condition (one-way ANOVA, or Log-Rank for survival curves), between nested conditions (Cox Proportional Hazards for survival curves), represented as *p < 0.05, **p < 0.01, ***p < 0.001, n.s. p > 0.05.

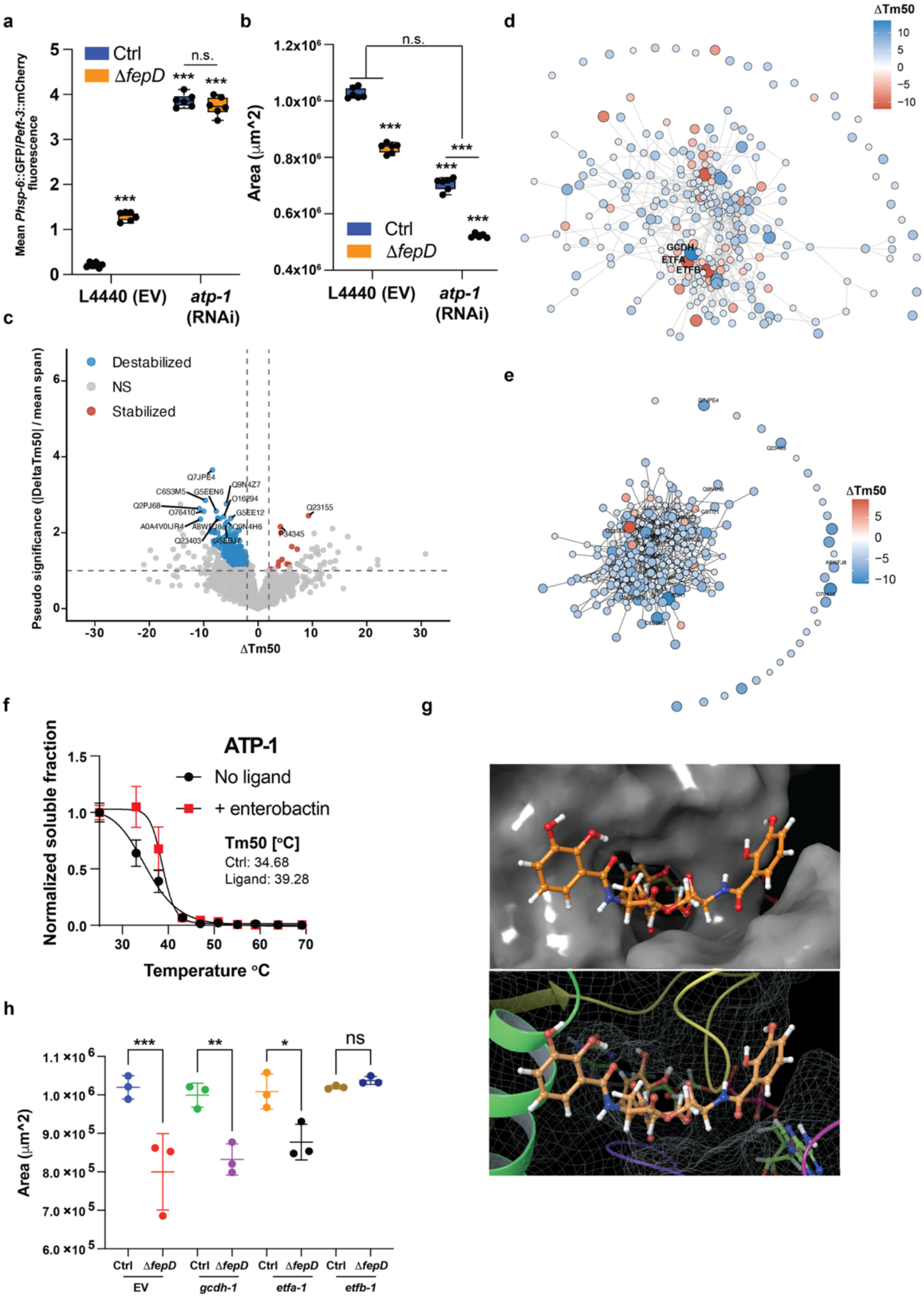


**Extended Data Figure 8. a,** Normalized *hsp-6* reporter intensity in N2 worms, quantified as the GFP/mCherry fluorescence ratio, following RNAi-mediated knockdown of *atp-1*. **b,** Changes in worm size, quantified by body area, in N2 worms following RNAi-mediated knockdown of *atp-1*. **c,** Proteome-wide protein stability changes in *C. elegans* following treatment with 100 μM DHB. Each dot represents one quantified protein. Protein stability is expressed as the change in melting temperature ($\Delta T_{m50}$), with positive values indicating stabilization (red) and negative values indicating destabilization (blue). Grey dots denote proteins with insufficient or non-canonical melting curve fits. **d,e,** STRING full-network analysis of significantly altered TPP hits in *C. elegans* following treatment with enterobactin (**d**) or DHB (**e**), using a confidence score threshold > 0.4. Nodes represent proteins with $|\Delta T_{m50}|$ > 2°C and pseudo-significance > 1, with node size proportional to $|\Delta T_{m50}|$ and node colour indicating stabilization (red) or destabilization (blue). **f,** Representative melting curves of ATP-1 in the presence or absence of 100 µM enterobactin. **g,** Predicted docking pose of enterobactin (orange) and FAD (green) within GCDH. **h,** Changes in worm size, quantified by body area, in wild-type worms following RNAi-mediated knockdown of *gcdh-1*, *etfa-1*, or *etfb-1*. Stats correspond to comparison against the indicated control (one-way ANOVA) and between nested conditions (two-way ANOVA), represented as *$p < 0.05$, **$p < 0.01$, ***$p < 0.001$, n.s. $p > 0.05$.

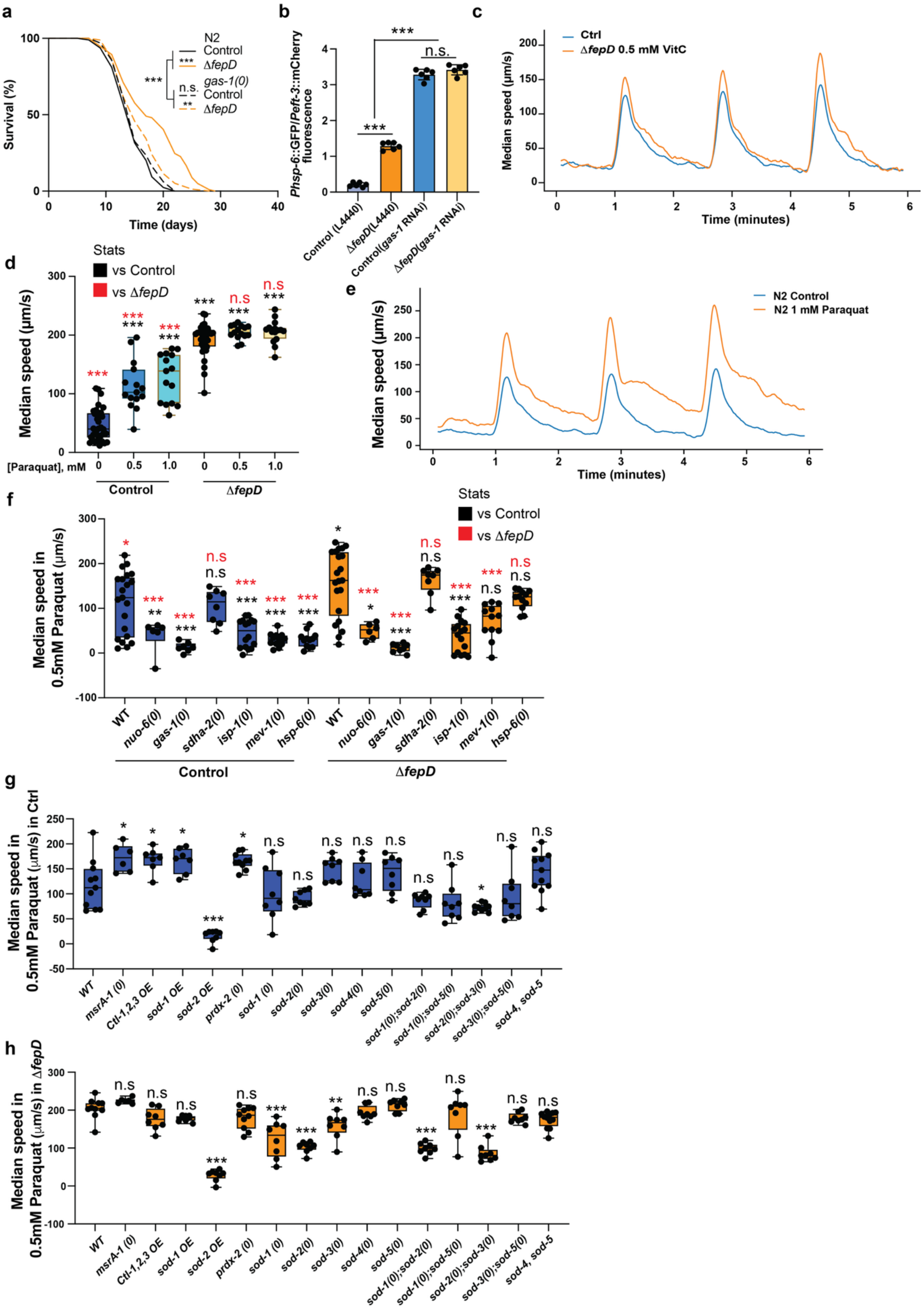


**Extended Data Figure 9. a,** Lifespan analysis of wild-type *C. elegans* (N2) and *gas-1* deletion mutants fed control *E. coli* BW25113 or Δ*fepD* bacteria (n=118-121, 2 independently run replicates) **b,** Normalized *hsp-6* reporter intensity in wild-type worms, quantified as the GFP/mCherry fluorescence ratio after feeding with the RNAi vector control or RNAi-mediated *gas-1* knockdown on BW25113 or Δ*fepD*. Each dot represents the average of eight worms (n=6 replicates). **c,** Time course of locomotor speed in response to blue light stimulation, indicated by the blue-shaded area, in worms fed control BW25113 or Δ*fepD* with 0.5 mM L-ascorbic acid supplementation. **d,** Median locomotor speed of worms fed control BW25113 or Δ*fepD*, with or without supplementation with 0.5 or 1 mM paraquat. **e,** Time course of locomotor speed in response to blue-light stimulation, indicated by the blue-shaded area, in worms fed control BW25113 with or without 1 mM paraquat supplementation. **f,** Median locomotor speed of wild-type worms and mitochondrial ETC mutants (*nuo-6, gas-1, sdha-2, isp-1, mev-1 and hsp-6*) fed control *E. coli* BW25113 or Δ*fepD*. **g,h,** Median locomotor speed of wild-type *C. elegans*(N2) worms, redox metabolism-related mutants (*msrA-1, prdx-2, sod-1, sod-2, sod-3, sod-4, sod-5, sod-1;sod-2, sod-1;sod-5, sod-2;sod-3, sod-3;sod-5 and sod-4;sod-5*), and antioxidant overexpression strains (*ctl-1; ctl-2; ctl-3, sod-1* and *sod-2* overexpression) fed either control *E. coli* BW25113 (**g**) or Δ*fepD* (**h**) with 0.5 mM paraquat supplementation. Statistics were performed against the control or Δ*fepD* mutant condition (one-way ANOVA, or Log-Rank for survival curves), between nested conditions (Cox Proportional Hazards for survival curves), represented as *p < 0.05, **p < 0.01, ***p < 0.001, n.s. p > 0.05.

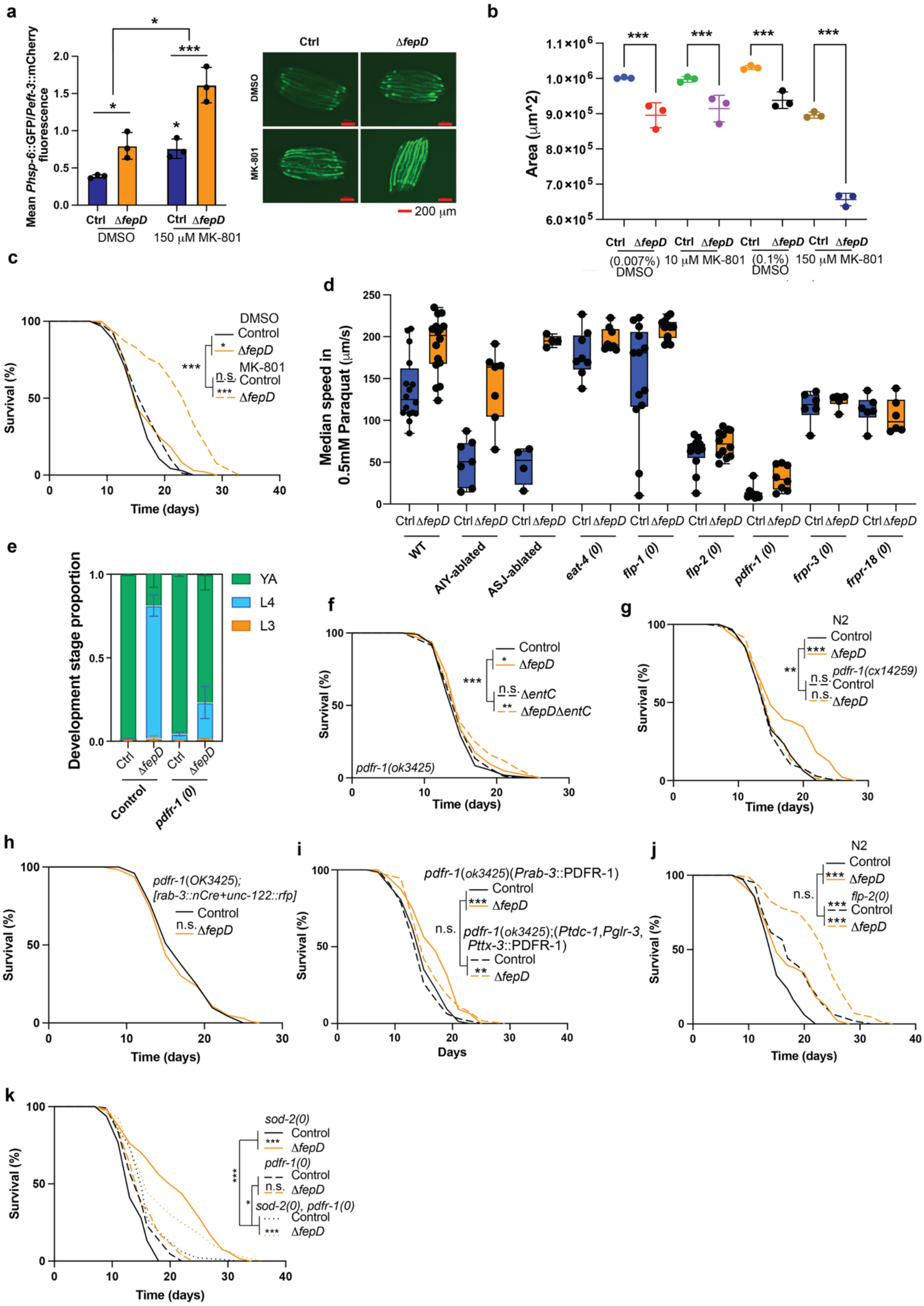

**Extended Data Figure 10. a,** Quantification and representative fluorescence images of normalized *hsp-6* reporter intensity in wild-type (N2) worms, measured as the GFP/mCherry fluorescence ratio after feeding with control *E. coli* BW25113 or Δ*fepD* bacteria, in the presence of 150 µM MK-801 or 0.1% DMSO vehicle control. **b,** Body size differences in wild-type *C. elegans* fed control BW25113 or Δ*fepD*

bacteria, supplemented with 10 μM or 150 μM MK-801, or the corresponding vehicle control. Body size was quantified by cross-sectional area, with each dot representing the average of 8 worms (n=3). **c,** Lifespan analysis of N2 worms fed control BW25113 or Δ*fepD* bacteria treated with 100 μM MK-801 (n=110-135, 2 independently run replicates). **d,** Median locomotor speed of wild-type worms or neuronal pathway-related mutant strains (*AIY*-ablated, *ASJ*-ablated, *eat-4, flp-1, flp-2, pdfr-1, frpr-3*, and *frpr-18*) fed control BW25113 or Δ*fepD* bacteria, in the presence of 0.5 mM paraquat. **e,** Developmental assay of wild-type and *pdfr-1* mutant worms fed control BW25113 or Δ*fepD* bacteria. **f,** Lifespan analysis of *pdfr-1* mutant worms fed control BW25113, Δ*fepD*, Δ*entC*, or Δ*fepD*Δ*entC* bacteria (n=119-121, 2 independently run replicates). **g,** Lifespan analysis of wild-type (N2) and *pdfr-1* mutant worms fed control BW25113 or Δ*fepD* bacteria (n=90-120, 2 independently run replicates). **h,** Lifespan analysis of neuronal Cre-modified *pdfr-1* mutant worms fed control BW25113 or Δ*fepD* (n=119-121, 2 independently run replicates). **i,** Lifespan analysis of pan-neuronal *pdfr-1* rescue worms, or *pdfr-1* rescue in *ttx-3-*, *glr-3-* or *tdc-1*-expressing neurons, fed control BW25113 or Δ*fepD* (n=125-140, 2 independently run replicates). **j,** Lifespan analysis of wild-type (N2) and *flp-2* mutant worms fed control BW25113 or Δ*fepD* (n=118-120, 2 independently run replicates). **k,** Lifespan analysis of *pdfr-1, sod-2,* and *pdfr-1; sod-2* double mutant worms fed control BW25113 or Δ*fepD* bacteria (n=121-134, 2 independently run replicates). Statistics were performed against the control or Δ*fepD* mutant condition (one-way ANOVA, or Log-Rank for survival curves), between nested conditions (Cox Proportional Hazards for survival curves), represented as *$p < 0.05$, **$p < 0.01$, ***$p < 0.001$, n.s. $p > 0.05$.
